# Aposematic color patterns are the dominant axis of phenotypic diversification in Nymphalid butterflies

**DOI:** 10.1101/2025.09.30.678834

**Authors:** Moritz D. Lürig, Leila T. Shirai, Luísa L. Mota, Keith Willmott, André V. L. Freitas, Arthur Porto

## Abstract

Butterfly wing patterns serve diverse roles in visual communication, from aposematic signaling and mimicry to mate attraction and camouflage. In brush-footed butterflies (Nymphalidae), this diversity can be traced to the wing pattern “ground plan” that generates phenotypes deterministic in origin yet highly multidimensional in form. Quantifying such complexity at scale has long been a challenge, limiting our understanding of how visual signals interact, constrain one another, and evolve. Here, we used computer vision to extract high-dimensional traits from standardized museum specimens and assembled the largest comparative dataset of wing color patterns to date, spanning over one third of all known Nymphalid species. We first tested whether chemically defended species occupy a distinct region in morphospace and then derived a quantitative score for aposematism from the principal color patterns associated with defense. Using this score, we examined whether aposematic signals are expressed consistently across wing surfaces and sexes, and whether their origins are linked to shifts in evolutionary rate. We found that the dominant axis of morphospace is defined by chromatic and achromatic contrast, along which defended and undefended species cluster. Validation with an expert-labeled moth dataset confirmed that this axis separates aposematic from non-aposematic phenotypes across Lepidoptera. Consistent with theory, strongly aposematic species showed greater visual similarity between dorsal and ventral surfaces, between sexes, and among individuals. Rate analyses further indicated that aposematic patterns evolved repeatedly and were associated with non-linear shifts in evolutionary tempo. Together, these results identify aposematism as the dominant organizing axis of wing color pattern evolution in Nymphalidae.

**Significance statement:** Butterflies are renowned for their striking diversity of wing patterns, including the warning colors that signal chemical defense to predators. Yet whether such warning patterns share common features across lineages has remained unclear. Here, we applied a metric computer vision model to more than 16,000 museum specimens spanning one third of all Nymphalid species, encoding their wing patterns into a common morphospace. Within this space, high-contrast aposematic patterns emerged as the dominant axis of diversification, explaining up to 20% of phenotypic variance. These signals were expressed consistently across wing surfaces, sexes, and individuals, and evolved repeatedly across the family. Our approach demonstrates how computer vision enables meaningful comparative analyses of complex patterns, revealing general principles of butterfly diversification.

## Introduction

Animal coloration plays diverse roles in visual signaling, influencing survival and reproduction through interactions with both conspecifics and predators (1). In butterflies, wing patterns have long served as a model for studying visual communication, with early recognition by Bates (2), Wallace (3) and Darwin (4), of the dual role in predator deterrence and mate attraction. Since then, extensive research has shown that the evolution of insect color vision (5) and adult diurnal activity in Lepidoptera (6) facilitated the use of color and color patterns for signaling (7). The stunning wing-pattern diversity of brush-footed butterflies (family Nymphalidae) manifested over 93 million years of evolution (8, 9) through permutations of a conserved developmental template dubbed the “ground plan” (10, 11), has been key to our understanding of how visual signaling evolves and drives diversification (12, 13). The permutational complexity of the ground plan has long hindered quantitative comparisons of pattern variation at scale. However, while recent applications of computer vision (CV) to images of pinned butterfly specimens have demonstrated how artificial intelligence can support comparative approaches (14–16), taxonomically broad mappings of biologically meaningful visual signals in butterflies still await initial explorations.

Aposematic color patterns, warning signals aimed at visual predators (primarily birds (17)), are among the most characteristic wing-phenotypes in Nymphalidae. Particularly well-studied are chemically defended species, which often advertise unpalatability through bright red or yellow hues arranged in patterns with high chromatic and achromatic contrast (18–21). Such patterns stand out against natural backgrounds (21, 22), and are thought to facilitate detectability and predator learning (23, 24). While specific color patterns are well documented and well-studied in a handful of taxa, it remains unknown whether there are shared, generalizable aspects of aposematic color patterns across the entire family, and how uniformly they are expressed. Signalling- and evolutionary theory predicts convergence on a single, consistent form driven by frequency-dependent selection (25–27), yet warning coloration is often remarkably diverse: among species, populations, individuals within a population, and even between dorsal and ventral wing surfaces of the same individual (28). Solving this “paradox” has been challenging due to the lack of standardized, large-scale quantifications of aposematic signals contained in wing phenotypes, also limiting our ability to reconstruct the origins and evolutionary history of warning coloration in Nymphalidae.

Here, we used a weakly supervised metric computer vision model (29, 30) to encode color patterns of standardized butterfly specimens from museum collections into a family-wide morphospace that encompasses approximately one third of all known species of Nymphalidae. Within this space, a dimension associated with conspicuous high-contrast color patterns emerges as the dominant axis of separation among chemically defended and undefended species, which accounts for up to 20% of the total phenotypic variance and exhibits strong phylogenetic structure. Leveraging this naturally emerging separation, we identified the color patterns associated with chemically defended species, hereafter referring to these patterns as “aposematic”, and tested for their prevalence across the family in a phylogenetically explicit framework. Specifically, we tested whether i) chemically defended species are phenotypically distinct from non-defended species along an axis of aposematic color patterns; ii) aposematic color patterns are consistently expressed across wing surfaces, sexes and individuals; and iii) aposematic color patterns have evolved repeatedly within Nymphalidae and alongside chemical defense. Our findings indicate that aposematic color patterns are a dominant axis of phenotypic differentiation, and were a major driver behind evolutionary diversification in Nymphalidae. Moreover, we demonstrate that computer vision features extracted from metric learning models, rather than hand-coded characters, allow for tractable comparative analyses of complex wing phenotypes, providing a framework to explain how aposematic phenotypes can vary in their expression yet remain consistently strong.

## Results and Discussion

We assembled an image dataset of Nymphalid butterflies that encompasses 16,725 museum specimens (2,543 species; Table 1), each photographed from both dorsal and ventral perspectives. From these images, we extracted image embeddings, high-dimensional vector-representations (N=768) of visual content, using a pretrained metric image encoder (see Methods section for more details). In short, metric learning learns a feature space in which distances encode biologically meaningful similarity among complex color patterns, enabling comparisons within a common morphospace (29, 30). To build this morphospace, reduce dimensionality and focus on the major axes of variation, we applied principal component analysis (PCA) on all embedding vectors (dorsal and ventral), which formed the basis for all downstream analyses. We then paired this phenotypic data with curated labels for chemical defense, which we assigned based on a literature review, and integrated sex, wing surface, and phylogenetic information to test our hypotheses. To do so, we used a combination of multivariate statistics, linear discriminant analysis, and phylogenetically informed models of trait evolution to characterize aposematic color patterns, quantify their consistency, and reconstruct their evolutionary history.

**Table 1.** Dataset overview. Our dataset encompassed images of 16,725 standardized specimens from 2543 species from museum collections, photographed from the dorsal and ventral side. We extracted image embeddings for all 2543 species, projected them into morphospace (Fig. 1), and used this information to calculate a score for aposematic color patterns (LD1). We collected information on chemical defense for 374 species contained in our dataset, and had sex information from the GBIF records for 1636 species, although this did not always include labels for both sexes per species (this was only the case for 1095 species). We obtained the phylogeny from Chazot et al. (9), which contained 1382 species from our dataset.

|  | Total |  | W. phylo. information |  |
| --- | --- | --- | --- | --- |
|  | Specimens | Species | Specimens | Species |
| Embeddings + LD1 | 16,725 | 2543 | 11887 | 1382 |
| Chemical defense labels | 4843 | 374 | 4698 | 355 |
| Sex labels | 8699 | 1636 | 6791 | 783 |
| Sex + chemical defense labels | 3484 | 369 | 3350 | 336 |
| Dorso-ventral similarity | - | 2543 | - | 1382 |
| Inter-sexual similarity | - | 1095 | - | 783 |
| Intra-specific similarity | - | 424 | - | 362 |

### Phenotypic differentiation along an aposematic axis

Using image embeddings, we identified aposematic color patterns as the dominant axis of phenotypic diversification in Nymphalidae. Across the family, chemically defended (see methods) species separated strongly from undefended species along a single axis in morphospace (P<0.001; Fig. 1; Table S1A), which explained between 14.4% (dorsal) and 20.7% (ventral) of the total variance (mvGLS; (31)). Specifically, chemically defended species occupy parts of morphospace marked by fine, vein-interspersed patching and high color (chromatic) and luminance (achromatic) contrast (Fig. S2) - phenotypes that were most consistently expressed in the subfamilies Danainae and Heliconiinae (Fig. 2). Subfamilies such as Satyrinae and Charaxinae, which contained mostly species with low chemical defense, occupied an area associated with overall greater color and pattern heterogeneity, encompassing both muted tones as well as variable colors, and large monochrome patches. Species with intermediate chemical defense, mostly from the Nymphalinae and Heliconiinae, were distributed diffusely in morphospace between the high- and low-defense clusters, yet also occupied a distinct region of morphospace characterized by fine-scale red to yellow dots and patches. Separation was more nuanced but also highly consistent when aggregated by sex (Fig. S3, Table S1A), with strongest separation in the female ventral surface (23.7% variance explained) and weakest in the male dorsal surface (14.3% variance explained). This consistency across wing surfaces and sexes, and the fact that this highly structured morphospace occupation emerged purely from unsupervised dimensionality reduction (i.e., PCA), suggests that chemical defense aligns with a major axis of evolutionary divergence in wing coloration.

**Figure 1.**
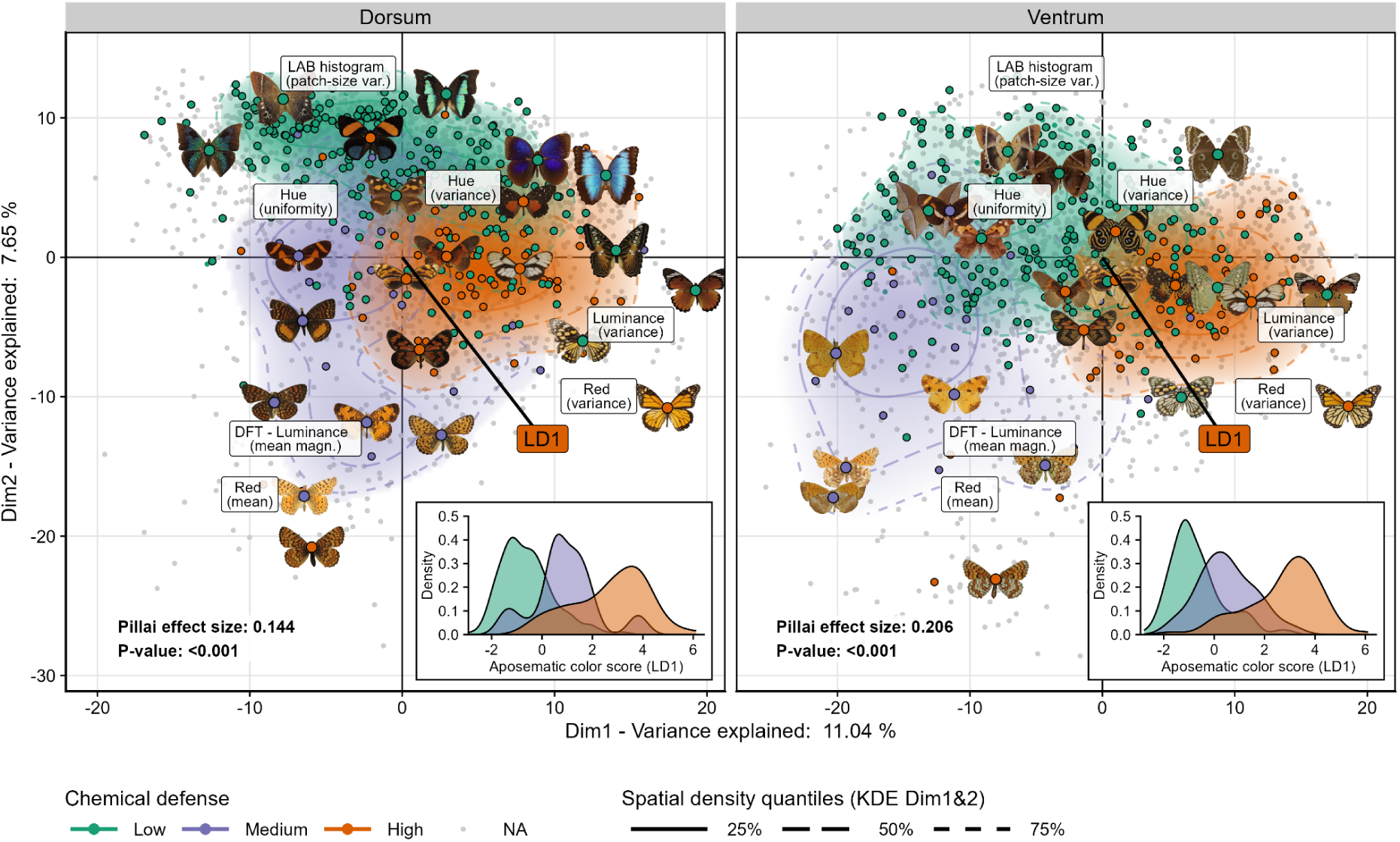
Morphospace of wing color patterns in Nymphalidae is structured by chemical defense. Morphospace of wing color patterns, constructed using a Principal Component Analysis (PCA) of embedding vectors from all specimens (N=16,725, Table 1). Each point is the species centroid in PC space (N=2,543), colored by chemical defense category; bold points indicate species from the labeled subset (no information for small gray points). Pictograms illustrate representative species associated with different regions of morphospace; color pattern features (e.g., luminance variance, hue uniformity) are projected as supplementary variables (with a slight jitter to avoid overplotting) to provide additional context for the axes (see Fig. S2 for more features and detailed explanations). Contour lines show 25%, 50%, and 75% spatial quantiles from a 2D kernel density estimate of the first two PCs; insets display the distribution of aposematic scores (LD1) across chemical defense categories. The vector labeled “LD1” indicates the alignment of LD classifier scores with phenotypic space (projected as a supplementary variable). Species categorized as low, medium, or high in chemical defense separate in multivariate PC-space, which we tested using a phylogenetically informed multivariate GLS (Table S1A; P<0.001 for both dorsal and ventral wing surface). Species in the low-chemical-defense group are characterized by large monochrome color patches and overall variable coloration, whereas species with high chemical defense and aposematic score (LD1) tended to have high luminance and chromatic contrast. Orthogonal to that axis were species with intermediate chemical defense, whose main characteristics are “checkerboard” patterns with high intensities of orange and red coloration (i.e., Fritillaries (47)).

**Figure 2.**
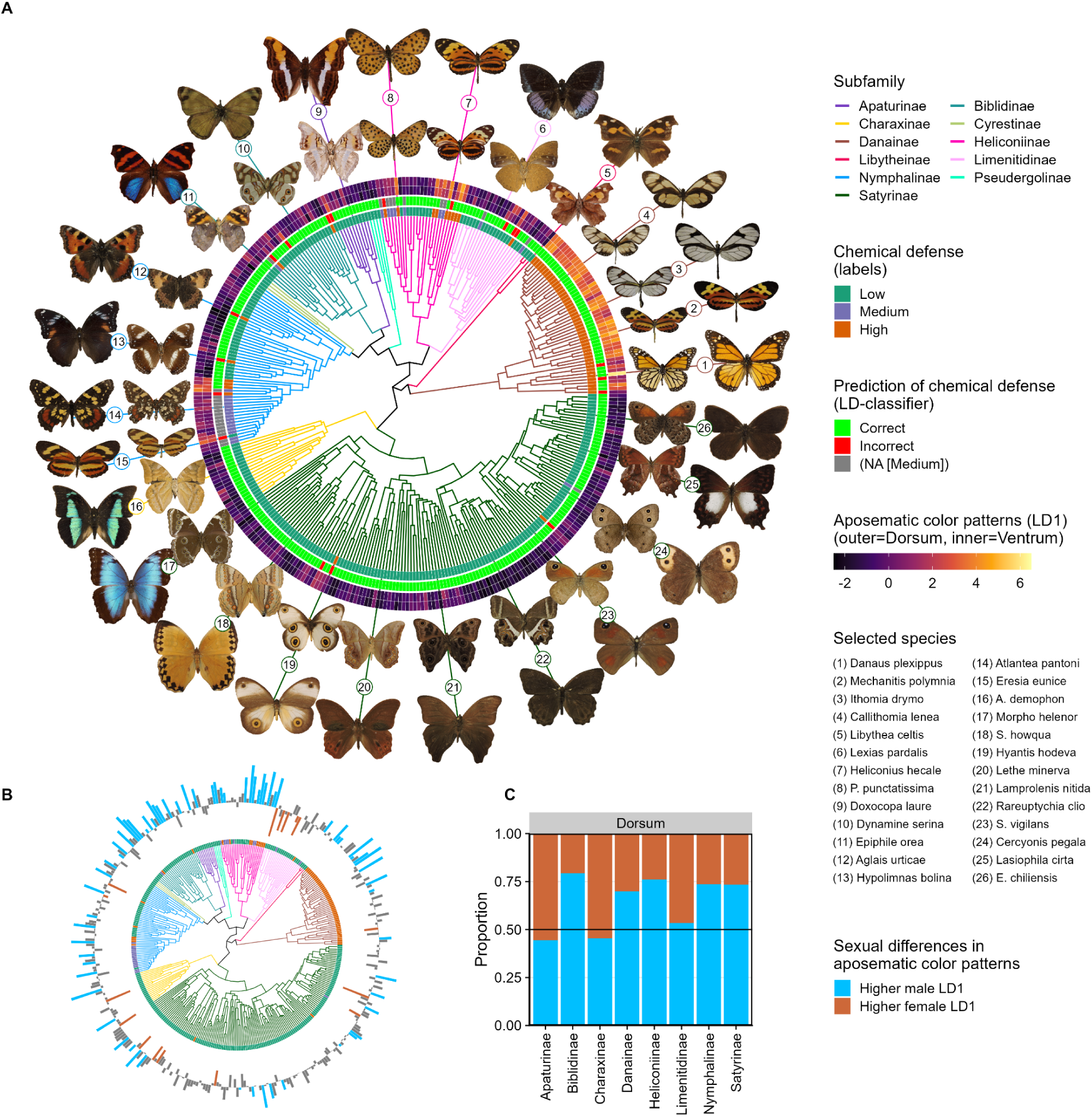
Aposematic color patterns have strong phylogenetic signal A: Ultrametric tree of all species with labels for chemical defense in the phylogeny (N=355; Table 1), with branch colors denoting subfamilies. Concentric rings show (from inside to outside): chemical defense categories based on literature reports (low, medium, high), predictions of chemical defense from the LD classifier (correct, incorrect, or NA for “medium”), and aposematic scores (LD1) for ventral (inner ring) and dorsal (outer ring) surfaces, ranging from low (purple) to high (yellow). Exemplary species are shown around the tree, corresponding to numbered labels. The distribution of aposematic scores demonstrates strong phylogenetic signal, with consistently high values concentrated in Heliconiinae and Danainae, but also independent origins in Nymphalinae and scattered cases within predominantly non-aposematic subfamilies such as Satyrinae and Limenitidinae. **B:** Phylogenetic distribution of species-level sex differences in aposematic score (male - female), with outward bars indicating higher scores for males and inward bars indicating higher scores for females. Coloration (blue=male, orange=female) indicates whether a species deviated from an expected null distribution of sex differences (Fig. S6) **C:** Proportion of species within each subfamily showing sex-biased aposematic scores on the dorsal surface (only species outside the null distribution; for ventral see Fig. S6), revealing pronounced male-bias in the strength of the aposematic signal in the five biggest subfamilies.

The observed separation between chemically defended and undefended species, as well as the color patterns we found to be associated with chemical defense, are consistent with existing theory and empirical work describing aposematic color patterns in butterflies (18, 19, 32), and the configuration of warning colors in general (20, 21, 27, 28). To extend this association beyond the labeled subset, validate it externally, and obtain a quantitative score for aposematism, we trained a linear discriminant (LD) model to isolate the primary axis of phenotypic separation associated with chemical defense in morphospace. For training, we used a subset of principal components (PCs) and only high- and low-defense species to reduce label ambiguity and maximize phenotypic contrast. Performance peaked with the first seven PCs (Fig. S4): 90.4% (dorsal) and 92.3% (ventral) for in-distribution species, and 91.4% (dorsal) and 86.4% (ventral) for out-of-distribution species (Table S2), indicating that LD1 captures a robust, generalizable signal. We then evaluated the model on an independent, expert-labeled moth dataset ((33); aposematism vs. camouflage). The LD model achieved 84.5% accuracy on the moth dataset (Table S2), and the two signaling strategies separated along the same morphospace axis as high- vs. low-defense butterflies (Fig. S5).

The finding that the observed axis of phenotypic differentiation extends beyond our focal clade supports the idea that aposematic color patterns are not only the dominant axis of phenotypic divergence in butterflies (34–36), but one that is visually conserved and evolutionarily meaningful across Lepidoptera (33), and potentially other taxa (5, 28). Although aposematism has long been understood as a powerful selective force that shapes prey appearance and predator behavior via signal-receiver coevolution (26, 37), only recently through digital imaging and CV have we been able to directly quantify its phenotypic aspects (32, 33). The repeated emergence of patchy, high-contrast wing patterns that we found in chemically defended lineages implies that aposematic signals may occupy only a relatively narrow region of accessible visual phenotypic space, putatively optimized for detection, memorability, and efficient learning by visual predators (26), or for exploiting their visual system (27). Indeed, the fact that we were able to achieve high classification accuracies on only seven phenotypic dimensions (i.e., the first seven PCs) indicates that there are only few generalizable aspects of aposematic color patterns, associated with patterns of high chromatic and achromatic contrast, but not specific color hues (Fig. 1; Fig. S2) (33). These pattern aspects, albeit manifested in diverse wing phenotypes, nevertheless define a consistent visual strategy that was measurable across the entire Nymphalid family, and beyond.

### Strong phylogenetic signal of aposematic color patterns

Aposematic score (LD1) exhibited strong phylogenetic signal (Blomberg’s k between 0.46 - 1.16; P=0.001; Table S1B), concentrated in Danainae and Heliconiinae, and, unexpectedly, in subsets of Nymphalinae (Fig. 2). In Danainae and Heliconiinae, aposematic color patterns have long been associated with mimicry rings that include many of the best-studied examples of Müllerian mimicry in butterflies (38, 39), such as *Mechanitis polymnia* (Disturbed Tigerwing; Fig. 2A, common names follow Geale (40) for South American species and various other sources for other regions) and other members of the tribe Ithomiini (e.g., *Ithomia drymo*, *Callithomia lenea*; Fig. 2A). *Danaus plexippus (*Monarch), from the tribe Danaini, showed the highest aposematic score (LD1) in our dataset; likely due to the combination of multiple conspicuous elements such as high-contrast vein- and dotted wing border patterns (41, 42). In Heliconiinae, strong aposematic signals were especially pronounced within the tribe Heliconiini, including some of the best studied aposematic butterflies, such as *Heliconius hecale* (Arrowhead Longwing; Fig. 2A) and *Heliconius erato* (Lovely Longwing). The Nymphalinae, in contrast, are not classical examples of aposematism or mimicry, yet our results show that a subset of species within this group, chiefly those belonging to the tribe Melitaeini (e.g., *Atlantea pantoni*), also exhibit relatively pronounced aposematic signals, consistent with our assessment of intermediate chemical defense. Across most subfamilies, when there were sexual differences in aposematic color patterns, males tended to have higher LD1 scores, particularly on the dorsum (Fig. S6; Table S3). These inter-sexual differences were most extreme in subfamilies with a strong aposematic signal, like Danainae and Heliconinae (Fig. 2B/C), but the male bias was most pronounced among Biblidinae and Charaxinae, hinting at a possible role of warning signals for mate attraction (43, 44)

We also found widespread genus-level consistency in the expression of aposematic color patterns: 86% of the genera containing more than one species in our dataset were overwhelmingly composed of either high- or low-defense species (i.e., >90% species assigned to the same group; Fig. S7). Among the largest genera in our dataset, color patterns in *Acraea* and *Heliconius* were classified as highly aposematic in over 90% (113/125) and 98% (52/53 species) of all cases, respectively, whereas *Charaxes* and *Limenitis* were classified as low-defense in over 97% (92/94 species) and 83% (50/60 species) of all cases, respectively. This genus-level cohesiveness suggests that visual defense strategies are often shared across closely related and often sympatric species (45). It is noteworthy, however, that training labels for ‘high-defense’ were concentrated in a few clades, which could potentially limit the classifier’s sensitivity to more divergent aposematic phenotypes. For example, species like *Aglais urticae* (Small Tortoiseshell; Fig. 2A) and *Hypolimnas bolina* (Great Eggfly), both of which are considered to be chemically defended and to have aposematic color patterns (46), were not classified as aposematic by our LD-classifier. Neither were the dotted and checkered patterns associated with a paraphyletic group referred to as “fritillaries” (i.e., “checkerboard” or “dice-box” butterflies) (47). In these genera, the scoring of aposematic color patterns was much more phylogenetically inconsistent, for instance, 58% in the genus *Boloria* (10/17 species) or 50% in the genus *Chlosyne* (12/24; Fig. S7).

### Consistency of visual signalling

Phenotypic similarity within species increased with aposematic signal strength along three axes (dorso-ventral, across sexes, and among individuals within a species after accounting for wing-surface and sex), across both aposematic and broader aspects of the phenotype (Fig. 3; Table 2A), and regardless of phylogenetic context (Fig. S8; Table S4). To quantify visual similarity, we used cosine similarity on two sets of PCs: one emphasizing aposematic variation (PC1-7; used for the LD classifier) and one representing overall wing color pattern variation (PC1–177, capturing 95% variance). In both cases, similarity increased significantly with aposematic signal strength, with slightly stronger effects for overall pattern similarity (Table 2A). Dorso-ventral similarity showed the strongest positive correlation with LD1 (Fig. 3A): in highly aposematic species like *D. plexippus* or *Oleria onega* (Split-spot Woodshade), dorsal and ventral wing patterns were nearly indistinguishable, whereas *Morpho helenor* (Common Morpho) or *Archaeoprepona demophon* (One-spotted Gladiator) featured brightly iridescent dorsal- and brownish muted ventral wing colors. This contrast likely reflects divergent evolutionary strategies: on the one hand, the aposematic signal in defended species (and their mimics) is maximized by being consistent, for instance, to stand out against different backgrounds (21, 22), and to reduce the number of signals predators have to learn (23, 48). On the other hand, lower dorso-ventral similarity likely reflects the partitioning of visual signals, such sexual signaling (49) or flash coloration dorsally (50), and camouflage (51, 52) or disruptive color patterns on the ventral surface (22, 53).

**Figure 3.**
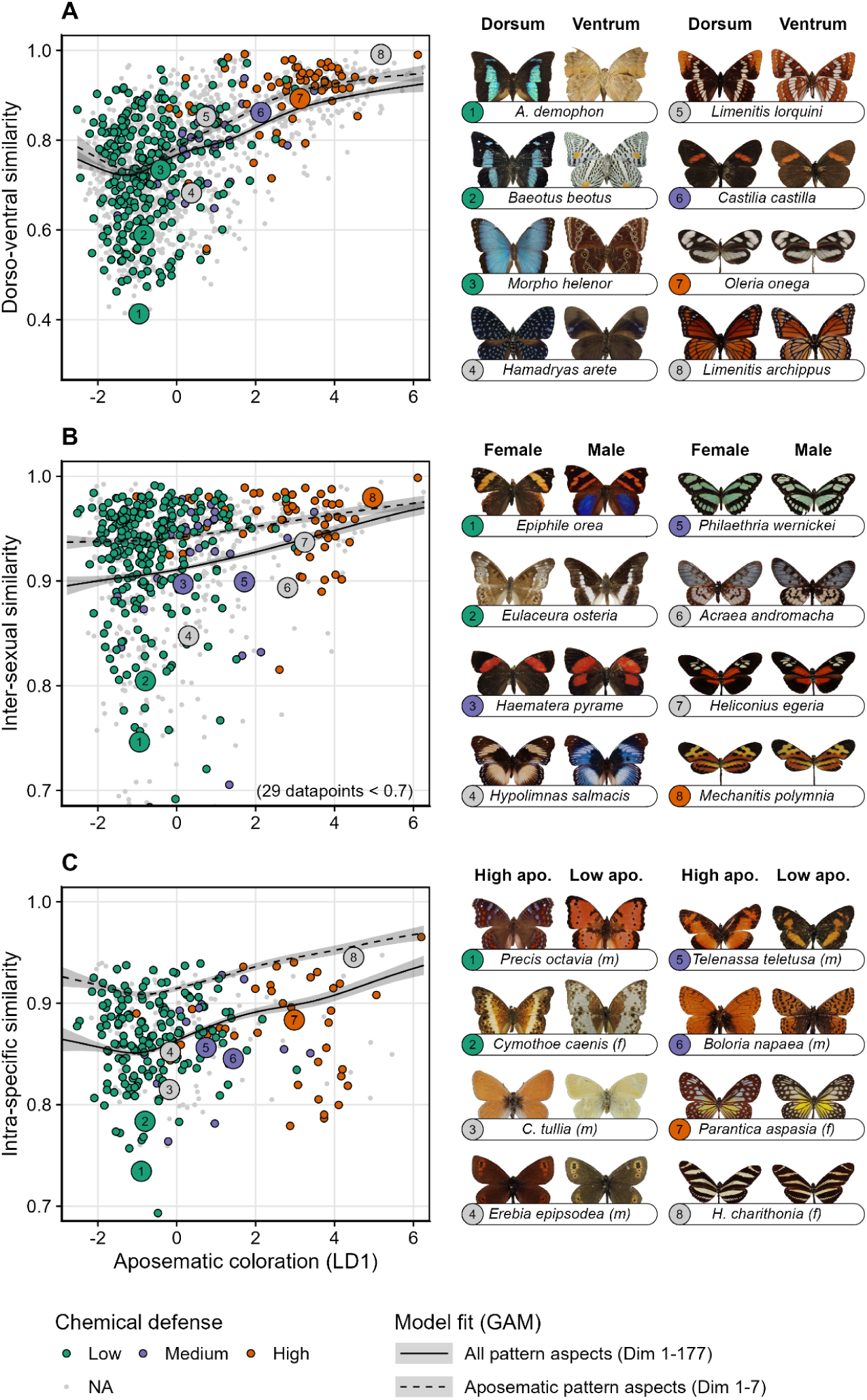
Aposematic color patterns are associated with reduced intraspecific variability. Each panel on the left shows the relationship between aposematic score (LD1) and different axes of intraspecific phenotypic similarity (A=dorso-ventral similarity [N=783], B=inter-sexual similarity [N=783], C=intra-specific similarity after accounting for wing surface and sex differences [N=362]). Points are colored by chemical defense, smaller gray dots denote species that were in the phylogeny but for which we did not have labels. Solid lines represent fits from Generalized Additive Models (GAM) based on all pattern aspects (PCs 1–177), while dashed lines represent GAM fits based only on aposematic aspects (PCs 1–7). The GAMs were fitted using all species from the phylogeny (i.e., colored and gray dots; see Fig. S5 and Table S4 for model fits using all available species), and the shown points indicate similarities calculated on all pattern aspects (i.e., corresponding to the solid line). Selected examples of species are shown in the image grids to the right (numbers correspond to the examples in the scatter plots). All shown fits are highly significant (see Table 2, and Fig. S5 for the partial fits of interaction terms).

**Table 2.** Results for aposematic score and phenotypic similarity A: Results for Generalized Additive Models (GAM) of phenotypic similarity (dorso-ventral, sexual, and intra-specific) over aposematic score (LD1). All shown models use the betar family (logit function), 10 knots per term, and include a phylogenetic penalty (knots=N species / 2). Shown columns are, from left to right: model names, response variables (All=similarity based on PC1-177, Aposematic=similarity based on PC1-7), terms for main effect (s(LD1)) and factor interaction smooths (s(LD1), …), phylogenetic penalty (s(species)), estimated degrees of freedom (Est. df.), reference degrees of freedom Ref. df.), Chi-squared statistic (Chi-sq.), and P-value (significant >=0.05 in bold). **B:** Result for GAM-regression of probabilities for a higher evolutionary rate over aposematic score (LD1), with the same model structure and columns as described for panel A.

| Model | Response | Terms | Est. df. | Ref. df. | Chi-sq. | P-value |
| --- | --- | --- | --- | --- | --- | --- |
| A |  |  |  |  |  |  |
| GAM 1a | D/V similarity<br>(All) | s(LD1) | 5.727 | 6.762 | 138 | < 0.001 |
|  |  | s(species) | 344.933 | 690 | 1967 | < 0.001 |
| GAM 1b | D/V similarity<br>(Aposematic) | s(LD1) | 5.269 | 6.29 | 105 | < 0.001 |
|  |  | s(species) | 320.541 | 690 | 1689 | < 0.001 |
| GAM 2a | Sexual<br>similarity<br>(All) | s(LD1) | 2.764 | 3.47 | 32.98 | < 0.001 |
|  |  | s(LD1,wing surface) | 2.001 | 2.002 | 41.03 | < 0.001 |
|  |  | s(species) | 246.314 | 390 | 1575.48 | < 0.001 |
| GAM 2b | Sexual<br>similarity<br>(Aposematic) | s(LD1) | 2.349 | 2.938 | 15.159 | 0.002 |
|  |  | s(LD1,wing surface) | 2.001 | 2.003 | 4.381 | 0.112 |
|  |  | s(species) | 189.323 | 390 | 1120.133 | < 0.001 |
| GAM 3a | Intra-specific<br>similarity<br>(All) | s(LD1) | 4.66 | 5.723 | 71.839 | < 0.001 |
|  |  | s(LD1,sex) | 2.001 | 2.002 | 6.039 | 0.049 |
|  |  | s(LD1,wing surface) | 3.317 | 3.897 | 15.361 | 0.004 |
|  |  | s(LD1,sex,wing surface) | 2.343 | 2.61 | 2.334 | 0.340 |
|  |  | s(species) | 124.021 | 180 | 500.443 | < 0.001 |
| GAM 3b | Intra-specific<br>similarity<br>(Aposematic) | s(LD1) | 4.21 | 5.194 | 45.502 | < 0.001 |
|  |  | s(LD1,sex) | 2 | 2.001 | 0.178 | 0.915 |
|  |  | s(LD1,wing surface) | 2.651 | 3.05 | 12.861 | 0.005 |
|  |  | s(LD1,sex,side) | 2.723 | 3.159 | 2.819 | 0.414 |
|  |  | s(species) | 113.097 | 180 | 411.81 | < 0.001 |
| B |  |  |  |  |  |  |
| GAM 4 | Evolutionary rate<br>(probability) | s(LD1) | 7.722 | 8.592 | 353.55 | < 0.001 |
|  |  | s(LD1,sex) | 4.479 | 5.368 | 67.01 | < 0.001 |
|  |  | s(LD1,wing surface) | 6.762 | 7.942 | 1038.93 | < 0.001 |
|  |  | s(LD1,sex,wing surface) | 3.513 | 4.173 | 54.54 | < 0.001 |
|  |  | s(species) | 147.817 | 161 | 3185.95 | < 0.001 |

Sexual similarity, quantified as the average cosine similarity between males and females, increased significantly with aposematic signal strength (Fig. 3B, Table 2A). Highly aposematic taxa, such as Clearwings (e.g., *Oleria onega, Methona themisto*), showed almost no sexual dimorphism in color patterns, while non-aposematic taxa such as Banners and Shoemakers (e.g., *Epiphile ora, Prepona hewitsonius*) expressed pronounced differences between sexes, involving brightly colored patches, UV-iridescence, or other ornamental patterns. For instance, iridescent dorsal patches in males of Great Eggfly (*H. bolina*; Fig. 2A) are actively used in courtship displays and thereby increase the chance of successful mating (54). Moreover, in *H. erato*, only females express UV photoreceptors, (55), and selection on pattern coloration, UV-reflectance, and efficacy as aposematic signal appear to jointly drive the evolution of male ornamentation (32, 56). Intraspecific similarity, which we quantified by averaging mean cosine similarities per sex and wing surface, also increased with aposematic signal strength (Fig. 3C, Table 2A). Sources of intraspecific variation include, for instance, seasonal polyphenisms, as observed in Gaudy Commodore (*Precis octavia*; Fig. 3C). To our knowledge, almost no studies have explicitly investigated intraspecific variability of aposematic color patterns in adult butterflies (but see (57), (58)), but similar findings in moths support the idea that warning signals are subject to stabilizing selection (33). This is consistent with predictions that aposematic species exhibit less within-species variation than cryptic taxa, likely because signal consistency makes predator learning more effective (23, 24).

Our findings confirm the longstanding hypothesis that aposematic signals are most effective when they are strong and consistent (25–27). High dorso-ventral similarity in species with aposematic color patterns has long been noted (59, 60), but not been systematically demonstrated across such a broad comparative scale, and, in the context of chemical defense (61), likely because of the challenges associated with the comparison of high-dimensional wing color pattern traits. Also, diversity in warning coloration has repeatedly been described as paradoxical (28, 43, 62), raising the question of how selection for signal consistency can be reconciled with extensive phenotypic variation within and among species. Our character-free approach provides a new perspective on this paradox, suggesting a possible explanation: although fine-scale phenotypic variation in pattern aspects like hue, saturation, patch shape or location is clearly present among aposematic species (Fig. 1, Fig. S2), the strength of the warning signal among chemically defended species remains comparable (e.g., compare pictures for #7 and #8 across all panels in Fig. 3). This observation suggests that the quality of aposematic signals in butterflies can largely be maintained during shifts of selective peaks (63), for instance, amid local adaptation (e.g., to different predators (64) or microhabitats (65)), sexual selection (e.g., via cooption of warning patterns as ornaments (32, 43) or vice versa (44)), or when changing seasonal forms (e.g., in response to temperature and humidity (66)).

### Evolutionary origin of aposematic color patterns

Ancestral state reconstruction indicated that the most recent common ancestor of Nymphalidae was non-aposematic, with warning coloration arising multiple times independently across the phylogeny (Fig. 4A; Fig. S9). In Danainae, a single origin of aposematic color patterns coincided with the acquisition of chemical defenses, producing a subfamily-wide association that has remained stable ever since (67, 68). In Heliconiinae, the transition occurred independently within the tribes Heliconiini, Argynnini and Acraeini, where aposematic wing color patterns are also tightly linked to chemical protection and mimicry (69, 70). In Nymphalinae, a separate origin was reconstructed for Melitaeini, consistent with prior evidence for the presence of chemical defense, and conspicuous patterning in *Euphydryas* and related taxa (71). Outside these groups, aposematic phenotypes emerged sporadically within clades (such as Biblidinae, Satyrinae, or Charaxinae) that are otherwise dominated by chemically undefended species (72, 73). These scattered cases did not always correspond to known chemical defense, suggesting alternative processes such as Batesian mimicry or convergence on high-contrast signals driven by other selective processes, such as sexual signalling (32, 43, 44). Taken together, these reconstructions highlight that aposematic color patterns in Nymphalidae do not have a single evolutionary transition, but instead multiple, independent gains that indicate convergence and repeated origins, with lineage-specific dynamics shaped by diverse ecological interactions and selective pressures (23, 26).

**Figure 4.**
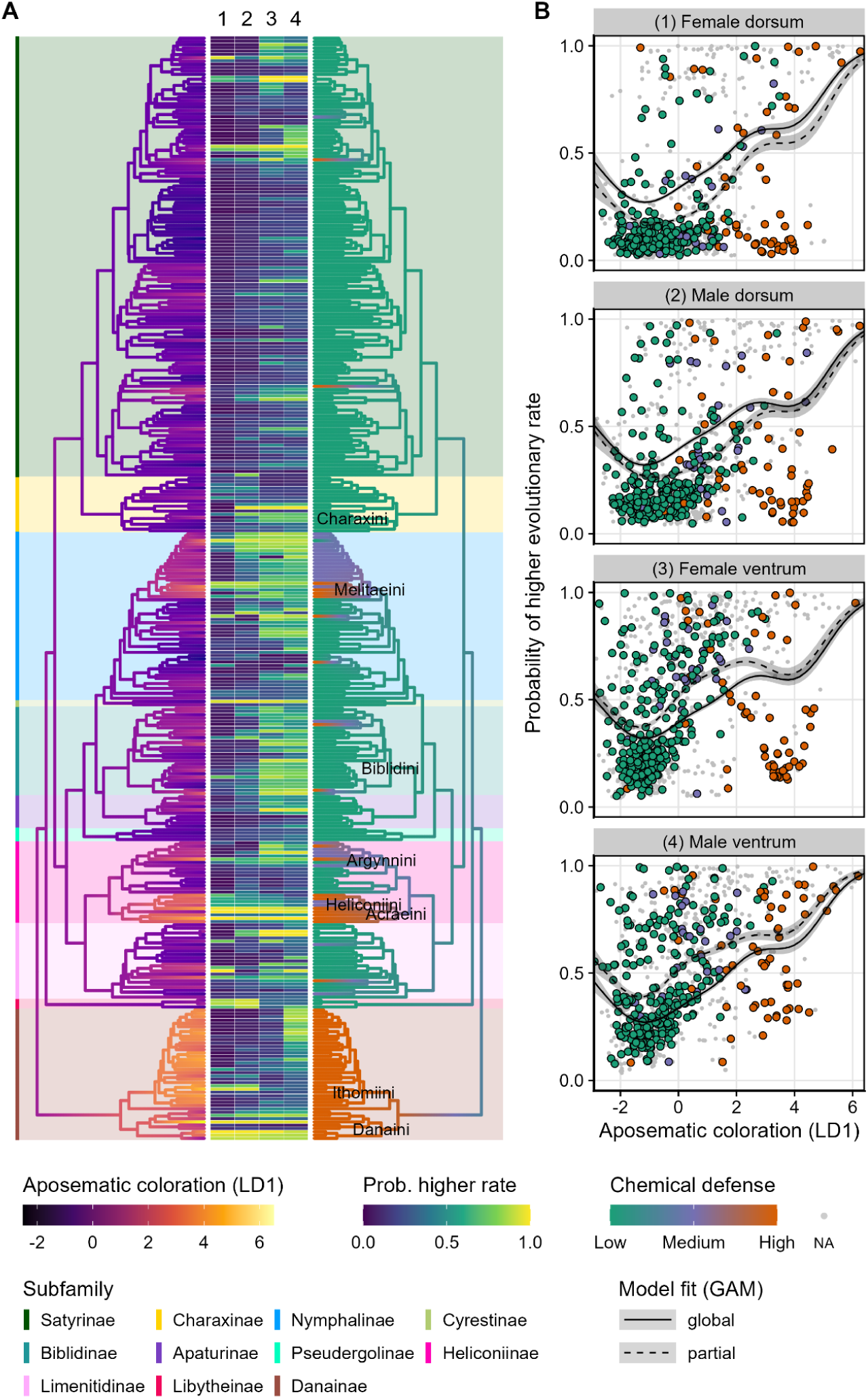
Repeated origins and elevated evolutionary rates of aposematic color patterns A: Composite panel shows, from left to right, ancestral state reconstruction of aposematic score (LD1) for all species with labels for chemical defense in the phylogeny (N=355; Table 1), estimated probabilities for a higher evolutionary rate for each wing-surface- and sex-context (numbers above the columns correspond to the facets in panel B), and ancestral state reconstruction for chemical defense. **B:** Each panel shows the relationship between the probability of high evolutionary rates and aposematic score (LD1). Points are colored by chemical defense; solid lines represent the global fit from a phylogenetically informed Generalized Additive Model (GAM; Table 2), dashed lines show the partial fits (i.e., the deviation for a specific subset). While there is a general positive relationship between elevated rates and higher aposematic scores, there are higher than expected rates at intermediate scores (resulting in a hump-shaped non-linear pattern). Moreover, evolutionary rates are higher on the ventral than on the dorsal wing side.

The lineage-specific clustering of aposematic color patterns was also reflected in our estimates of its evolutionary tempo. On the one hand, rates of evolution were higher on ventral than on dorsal wing surfaces (Fig. 4B; Table 2B; Fig. S9), which highlights the possibility for different selection regimes: for instance, stabilizing selection for sexual signalling dorsally (32, 43), and selection for either aposematism or camouflage ventrally (24). On the other hand, rates increased with the strength of the aposematic signal, which reflects the evolutionary gains of aposematism from a non-aposematic ancestor (Fig. 4A). Here, the Ithomiini represent a noteworthy exception: despite having the highest aposematic scores, species in this tribe were associated with relatively low evolutionary rates, in contrast to other typically highly aposematic tribes such as Heliconiini, Argynnini, Melitaeini, and Danaini. This indicates that the transition to aposematism in Ithomiini occurred relatively early, likely preceding the rapid diversification in this group that followed major geographic and ecological transitions in their Neotropical habitats during the Miocene (74), and a strong aposematic signal has remained consistent, despite the evolution of wing transparency (34). Other mostly aposematic lineages, like Heliconiini, do contain more frequent shifts in aposematic signal strength, and hence are associated with higher rates. We also found localized patches of unexpectedly high rates in clades that are otherwise characterized by low or intermediate aposematic scores, for example, within the tribes Biblidini and Charaxini, but also scattered across the tribe Satyrini at the species level, echoing the multiple, independent gains of aposematic color patterns identified in our ancestral reconstruction. Interestingly, we found a localized peak of evolutionary rates at intermediate aposematic signal, which was particularly pronounced for ventral wings of females (Fig. 4B). A possible explanation may be that evolutionary transitions between different signalling strategies are costly, and the resulting intermediate wing-phenotypes are maladaptive, reflecting instability while crossing fitness valleys between adaptive peaks (23, 26).

### Evidence for mimicry of aposematic color patterns

Although we did not intentionally pursue investigations regarding Batesian mimicry of aposematism, where undefended species mimic the appearance of defended species (75), our classification via the LD model revealed several cases where the model failed to predict the correct class, including species already recognized as mimics. In total, 36 out of 355 labeled species had a divergent class assignment, of which 19 low-chemical-defense were classified as “high” (Table S5). Among these species, the most striking case was *Pseudacraea poggei* (False Monarch), which is a well-known Batesian mimic of danaine butterflies (68). Moreover, the whole genus *Eresia* and other related Melitaeini (e.g. species of the genus *Castilia*) are known to form complex mimicry rings with Danainae and Heliconiinae (e.g., *Eresia lansdorfi* - False Erato), and the similarity of misclassified *Eresia eunice* (Tiger Crescent) to *H. hecale* (Arrowhead Longwing) supports the idea that selection from visual predators drives convergence in aposematic color patterns across distantly related taxa (Fig. 2). Surprisingly, *Consul fabius* (Tiger Leafwing), another member of the “tiger” mimicry ring, was not classified as aposematic at the species level, and only a few specimens were classified as aposematic on the dorsum (Fig. S7). Indeed, in several low-defense species aposematism was assigned to only some specimens, one wing-surface, or in one sex, such as the tiger-pattered females of the satyrine *Eteone tisiphone* (Table S5, Table S6). However, there were no systematic sex differences in these misclassifications, and thus no support for female-limited mimicry (Fig. S10, Table S5, Table S6), which is widespread in other butterfly families such as Papilionidae (3, 62, 76) and thought to reflect sex-specific predation risk. Still, the lack of sex data in many records currently limits definite conclusions on this matter.

A subset of other misclassifications involved species with visual resemblance to well-defended groups but for which little or no corroborating chemical or behavioral evidence exists. *Pseudohaetera hypaesia* (One-lined Sierra-Phantom), for instance, has highly transparent wings with subtle orange and brown markings that are strikingly similar to the pattern-arrangement of clearwings (Ithomiini). There is no evidence for chemical defense in this species, and it is currently unclear whether its visual similarity to ithomine species is the result of mimicry, or the shared evolutionary benefit of transparency in shady forest understory (34, 77). *Sephisa dichroa* (Western Courtier), in the subfamily Apaturinae, displays a high-contrast combination of black with contrasting orange and white patches, similar to certain fritillary specie. While its conspicuous dorsal patterning suggests the potential for visual mimicry, there are no records of it participating in established mimicry rings, and its host plants (mostly oaks) provide no known defensive compounds. *Vanessula milca* (Lady’s Maid) is a distinctive member of the Nymphalinae with reddish-brown wings and prominent white spots. Early literature speculated that it might mimic unpalatable Acraeini such as *Telchinia* (78), but these ideas have not been tested experimentally, and its larval host remains unknown. In each of these cases, the visual similarity to aposematic models is compelling enough to explain their misclassification by the LD-based scoring, yet the underlying ecological and chemical context is poorly resolved. Without targeted studies of host use, toxin sequestration, and predator responses, it remains unclear whether these patterns function as true warning signals, incidental resemblances, or adaptations for other selective pressures such as mate recognition or predator evasion.

### Conclusions and Outlook

Our study demonstrates that computer-vision-derived visual features offer a powerful and assumption-light framework for quantifying and comparing complex phenotypes across taxa. By decomposing variation into principal dimensions, we were able to identify the strong and recurring signal of aposematic color patterns as a dominant axis in phenotypic diversification in Nymphalidae. This aposematic signal not only distinguishes chemically defended species with high accuracy, but also shows remarkable consistency across wing surfaces and sexes, suggesting strong stabilizing selection for within-species consistency. Additionally, we found that although different mimicry complexes may differ sharply in color pattern, they are similar with respect to the aposematic strength of their color patterns. This supports the broader notion that variation within aposematic systems is not paradoxical, but bounded by perceptual constraints that are shaped by visual systems of animal predators. Signal consistency may therefore represent a fundamental principle not only in butterfly evolution, but potentially across aposematic organisms more broadly. Given the central role of birds as visual predators of butterflies and moths, future work should aim to place putatively aposematic phenotypes in predator-relevant perceptual space, for instance, by incorporating avian spectral sensitivity, ultraviolet imaging, and ideally testing predictions in behavioral assays.

## Materials and Methods

For an overview of the data-collection and analytical pipeline see Figure S1.

### Image acquisition and segmentation

We compiled an image dataset of Nymphalidae based on occurrence records from GBIF and iDigBio that met three criteria: they were based on physical specimens from museum collections, included at least one associated image, and the species was listed as “accepted” in GBIF’s taxonomy backbone. On October 14th, 2024, we queried GBIF with the above mentioned conditions, and downloaded a CSV file with which contained 369,978 observation records and external download links to associated images. We then downloaded all available images at their native resolution onto a local workstation using a Python-based protocol, which also avoided the creation of duplicates by calculating hash-based image checksums (MD5-protocol), resulting in 359,804 unique images. Based on the hash we also excluded images associated with multiple species (e.g., whole drawer images), as we would not be able to definitely associate them with their corresponding observation record.

We further supplemented this dataset with images of every species from Wahlberg et al. (8), including geographical races or subspecies, and sexual and seasonal dimorphism, based on a comprehensive taxonomic literature assessment for each species or, if not found, the genus that species represented. In total, we imaged 2,687 specimens belonging to 399 species with males and females represented, from both dorsal and ventral side (i.e., 5,374 images). The majority of specimens were photographed at the Biological Diversity Museum at the State University of Campinas (Unicamp), the Coleção Entomológica Padre Jesus Santiago Moure at the Federal University of Paraná (DZUP), and the McGuire Center for Lepidoptera and Biodiversity at the Florida Museum of Natural History (MGCL). Additional species were found at the Zoology Museum of the São Paulo University (MZUSP), the American Museum of Natural History (AMNH), and the Smithsonian National Museum of Natural History (NMNH) entomological collections.

We segmented the butterfly specimens from all images using GroundedSAM (Ren et al. 2024), a foundational computer vision model that segments objects with pixel-level accuracy based on text prompts (we used the prompt “butterfly”). We saved all segmented specimens as 3-channel RGB images, with the background masked in black. After segmentation, we were able to exclude images containing more than two specimens, which was the maximum we could safely associate with either dorsal or ventral side via a trained classification model (see below). After all these steps, we were left with a total of 529,887 segmented butterfly masks from 4,104 species. We use the term species for consistency throughout the paper, but the taxonomic status of some of these butterfly taxa may not be clear.

### Dataset assembly (embeddings, classification and labels)

We implemented a processing pipeline that relied on image embeddings, i.e., vector-based representations of two-dimensional visual content. Image embeddings would represent the segmented butterfly masks in a format suitable for quantitative analysis, and also allow us to remove erroneous masks from the dataset (e.g., incomplete or inaccurate detections, paper labels, rulers, etc.). To generate embeddings, we used UNICOM (30), an image encoder model pre-trained on 400 million images, that converts each image into an embedding vector with 768 dimensions. First, we projected the vectors into a common space via t-Stochastic Neighbor Embedding (tSNE), to identify clusters for further downstream processing, cleaning and analysis. In tSNE-based embedding space, objects that are perceptually more similar to each other are grouped closer together and possibly form clusters (e.g., all striped butterflies, all brown butterflies, all paper labels, etc.), whereas objects that are more dissimilar will be further apart. Then, using interactive scatterplots created with the *bokeh* library (79), we selected data-points that belong to clusters of erroneous segmentations, as well as clusters of accurately segmented butterflies. We selected both approx. 21,000 clean butterfly masks and “junk” masks to train a binary classifier model using the YOLO framework (80), with a 0.8/0.1/0.1 train/validation/test split. The resulting model achieved 99.7% accuracy on the test set, and we applied it to all masks to retain only accurately segmented butterflies (237,591 masks). We further refined the dataset by removing outliers based on the contour coordinates (i.e., non-butterfly shaped masks) to remove masks missed by the junk-classifier (230,689 masks).

We used information from the downloaded records, combining both metadata fields and parsed URL content, to identify whether each image showed a dorsal or ventral perspective. Based on this classification, we compiled a training dataset containing approximately 8,400 images for each side, and trained a binary classification model in the YOLO framework with a 0.8/0.1/0.1 train/validation/test split. The resulting model achieved 99.6% accuracy on the test set, and we applied to all masks to classify them into dorsal and ventral perspective, resulting in 189,541 dorsal and 41,148 ventral images. At this point we also removed all records with incomplete or placeholder species names (e.g., genus-only classification, or sp. and spp., etc.), as well as obvious misclassifications such as the moth genus *Acidalia* (incorrectly listed as a nymphalid butterfly). Selecting from this pool of images, our final dataset included only specimens with exactly two images: one dorsal and one ventral, resulting in 33,450 masks representing 16,729 specimens from 2543 species.

We then assigned the subfamily to each of these species based on Wahlberg et al. (8), supplemented by information from the online databases *Butterflies of America* (81), *Lepidoptera and Some Other Life Forms* (82), or the original species descriptions where necessary. Moreover, for 8,699 specimens from 1,636 species, we obtained sex labels from GBIF and iDigBio metadata: 5,107 males and 3,592 females. No individual species showed a significant deviation from a 1:1 sex ratio after FDR-corrected binomial tests. However, a one-sample *t*-test on the overall male–female difference per species revealed a consistent male bias (mean = 0.93; 95% CI: 0.82–1.03; *t* = 17.95, *df* = 1,635; *p* <0.001). On average, species were represented by 3.3 males and 3.0 females, indicating a moderate but widespread overrepresentation of males. Overall, we expect that this imbalance does not affect our analyses, which focus on variation across species and sexes, and assume broad representation of both sexes across taxa, which our dataset provides.

### Literature survey on chemical defense

We conducted a literature survey to assign the probability that a certain taxon is unpalatable due to chemical defense for all 399 species in the phylogeny of Nymphalidae from Wahlberg et al. (8), which was the most current reference at the start of our work (our comparative analyses use the more recent phylogeny by Chazot et al. (9) - see *Statistical Analysis*). To assess unpalatability via chemical defense across these species, we used two complementary approaches: first, we reviewed existing meta-analyses and review-articles for evidence such as the presence of defensive chemicals in adults or documented unpalatability to predators. Second, we performed a structured literature search using Google Scholar, querying each subfamily, tribe, and genus in our dataset with the terms “palatable” (which also captured “unpalatable” and “palatability”) or “toxic.” For each taxon, we examined the first five pages of results per term, totaling up to ten pages per group, which yielded information for a total of 399 species from the phylogeny, 356 of which were also included in our image dataset.

Based on the collected evidence, we assigned each species one of three levels: low/untested, intermediate, or high. These categories reflect inferred probabilities of a species being unpalatable due to chemical defense from information derived from the literature, rather than confirmed or quantitative measurements of chemical defense (e.g., “highly” or “mildly” unpalatable). Most species have not been formally tested, and even for those that have, defense levels may vary geographically and within populations (83, 84). Additionally, predator responses to unpalatable butterflies can differ across species, individuals, and ecological contexts such as satiation (85). We excluded anecdotal claims and considered tests with ants to be weak evidence, given that ants are not visually oriented predators, although groups unpalatable to birds are often also unpalatable to ants (86, 87). Despite taxonomic changes over time, we believe our combined survey strategies captured most of the relevant literature and provide a robust overview of palatability patterns in Nymphalidae.

Species assigned to “high” included those with direct evidence of unpalatability or chemical protection, whereas the “low/untested” category included species shown to be palatable or lacking chemical defense. We categorized the remaining species based on patterns observed in other members of the same tribe: all species in the tribes Ithomiini, Danaini, Acraeini, and Heliconiini were labeled as “high”, given the well-documented prevalence of chemical defense in these groups (85, 88, 89). Exceptions include *Dryas* and *Philaethria*, which were shown to be palatable in predation trials (85, 90) but still assigned to the “intermediate” category due to evidence of weak chemical protection (91). We labeled untested species of Argynnini as “intermediate” because tested members are unpalatable and other Heliconiinae tribes are widely defended. We categorized species from Vagrantini as “low/untested” due to a lack of evidence, and Melitaeini to the “intermediate” category, as most tested species are unpalatable, including one that feeds on non-iridoid host plants (71, 92). We categorized all remaining untested species from other tribes as “low/untested.”

### Statistical analysis

#### Overview

We conducted all statistical analyses using R v4.4.2 (93). Our overarching goal was to investigate how wing color pattern variation relates to ecological and biological factors - chemical defense in particular, but also wing surface, sex, and phylogenetic history. To this end, we leveraged our curated dataset of specimen-specific image embeddings, as well as labels for wing surface and sex, and integrated it with species-level information for chemical defense, subfamily, and phylogenetic relatedness (see Table 1 for sample sizes). The phylogenetic tree we used was based on Chazot et al. (9) and provided information for 1,382 of the species in our dataset. Prior to testing our hypotheses, we reduced the dimensionality of the image embedding vectors using a single principal component analysis (PCA) that projected dorsal and ventral wing surfaces into the same coordinate space. Based on this morphospace we calculated a series of univariate traits that we would then use to test out hypotheses: specimen specific scores for the intensity of the aposematic signal and dorso-ventral similarity, as well as species-specific metrics for inter-sexual similarity, and intra-specific similarity after accounting for differences between wing-surfaces and sexes.

#### Quantifying aposematic color patterns

To quantify the intensity of aposematic color patterns, we trained a linear discriminant (LD) classifier to define a single univariate axis between species from the labeled subset with either “low” or “high” probability of chemical defense, but excluding those classified as “intermediate” to ensure a stronger contrast. Prior to training the classifier, we removed outliers from the training set using Gaussian mixture models (GMMs) fitted separately for “low” and “high” classes. For each class, we modeled the multivariate distribution of PCA coordinates using a single-component GMM and computed Mahalanobis distances from the class centroid (across dorsum and ventrum), excluding specimens with distances exceeding the 95% chi-squared threshold. We then trained the LD using an 80/20% split stratified by chemical defense on the PCs aggregated by species and wing surface (see Table S2 for sample sizes). To determine the optimal number of components we trained across a range of 1-100 PCs and then evaluate performance on four datasets: the training set, an in-distribution test set (the 20% split not used for training), and an out-of-distribution test set (the species excluded via GMM). To further validate this approach, and to examine whether the axis we’re isolating from morphospace aligns with color patterns that are broadly considered aposematic, we used a cross-taxon set of moths species from multiple families as external validation of our approach (33), which were labelled independently by experts through literature research as having either “aposematic” or “camouflage” coloration (Fig. S5). We processed the moth images, which featured only the dorsal perspective, in the same way as the butterfly images (see section “Dataset assembly”), and then projected the resulting embeddings into the butterfly PCA. Inspection of the results indicated that a model with 7 PCs performed best across all datasets and evaluation metrics (Fig. S4). After training, we used the LD model to assign an LD score to the dorsal and ventral side of every specimen from our dataset. We hereafter refer to this axis (LD1) that represents “aposematic color patterns,” with the understanding that, although we did not directly test for aposematism, it likely captures wing phenotypes commonly associated with warning coloration.

#### Similarity metrics

Next, we calculated similarity metrics using two sets of principal components (PCs): the first 177 PCs, which together explained up to 95% of the variance and captured overall color pattern variation; and the first 7 PCs used in the LD classifier, which specifically captured the aposematic signal. We decided to use the PC-vectors from dimensionality reduction rather than the raw embedding vectors to exclude irrelevant pixel-level variation (e.g., slight misalignments or lighting artifacts) and retain only the most biologically meaningful structure in the data. We used cosine similarity to quantify the similarity between dorsal and ventral views of each specimen (also aggregated by sex within species), and sexual similarity at the species level (separately for dorsal and ventral views), considering only species with at least one male and one female specimen. To quantify intraspecific similarity, we included only species with at least four specimens per sex. We then calculated the mean cosine similarity among all individuals within each sex and wing surface (i.e., female-dorsal, female-ventral, male-dorsal, male-ventral), and averaged across these four values.

#### Hypothesis testing

We tested our first hypothesis using phylogenetically informed multivariate analysis of variance, which we implemented using the mvMORPH package (31). For the model we calculated the centroids for each of the 355 species present in the phylogeny, using the first 177 PCs, and then fitted seven separate models using the *mvGLS* function: a single model using species centroids across all wing-surfaces and sexes, one set of two, each using the species means for dorsum and ventrum, respectively (Fig. 1), and one set of four, with all combinations of sex and wing surface (Fig S2; Table S1A). Each model had the PC matrix as the dependent variable, and the vector of labels for chemical defense as the independent variable. We fit all models with the Pillai test statistic and 1000 iterations. To complement the multivariate tests we plotted the first two Dimensions of the PCA (Fig. 1; Fig. S3), where we show the centroids for all species (including those without phylogenetic information), and the distribution of chemical defense (using 2D kernel density estimates). Moreover, we used the *phylosig* function to calculate Bloomberg’s k (94) and to test for statistical significance of the phylogenetic signal (Table S1B).

To test our second hypothesis, we fitted a series of generalized additive models (GAMs) using the *mgcv* package (95), with aposematic score (LD1) as the predictor, and each of the three similarity metrics as dependent variables (Fig. 3). In total, we fitted six models: three each with similarity metrics derived from 177 PCs (i.e., all phenotypic aspects) and three with seven PCs (aposematic aspects only). All six models use the betar family (logit function), 10 knots per term, and include factor smooth interaction terms for sex or wing surface in a hierarchical fashion where appropriate (see Table 2 for details), fitting both global (Fig. 3; across all contrasts) and partial smooths (Fig. S8). To account for phylogenetic relationships, we fit each model with a Markov random field (MRF) penalty matrix derived from the phylogeny (95), using a number of knots set to half the number of species. We assessed significance using approximate χ²-tests (Table 2A). Additionally, we fitted these models without a phylogenetic term to include all species, also those not present in the phylogeny (Fig. S8; Table S4).

We tested our third hypothesis using ancestral state reconstruction under maximum likelihood, and by fitting a multi-rate discrete character dependent model for continuous trait evolution of aposematic color patterns, both using the phytools package (96). First, we implemented ancestral state reconstruction for aposematic score (LD1) and for chemical defense (converted to a numerical score from 1 [low] to 3 [high]) using phytool’s *fastAnc* function, including only the species with labels for chemical defense (N=355, Table 1) for direct visual comparison (Fig. 4A). We then fit four (one for each combination of sex and wing surface) hidden rate models (HRM) under an equal-rates discrete transition assumption and with a binary hidden state across 100 levels using the *fitmultiBM* function, using all species with sex information in the phylogenetic tree (N=783; Fig. S9). Using the resulting HRM fits, we then computed marginal ancestral states to estimate the posterior probability of each species evolving under the higher-rate regime. Finally, to test whether these probabilities increased with the strength of aposematic signal, we fit a single GAM with the probability of being in the higher-rate regime as the dependent variable, and aposematic score (LD1) as the independent variable. The model uses the betar family (logit function), 10 knots per term, and includes a factor smooth interaction terms for sex or wing surface to fit global and partial smooths (Fig. 4B), as well as an MRF-based phylogenetic penalty term, number of knots set to half the number of species. We assessed significance using approximate χ²-test (Table 2B).

## Acknowledgements

We thank the following curators and technicians for their outstanding support at museum collections: Artur Nishibe Furegatti, Jean Fanton, and Michela Borges (Unicamp); Andrei Sourakov (MGCL); Mirna Casagrande and Olaf Mielke (DZUP); Renato O. da Silva and Marcelo Duarte (MZUSP); David Grimaldi (AMNH); and Bob Robbins (NMNH). A.V.L.F. thanks the Conselho Nacional de Desenvolvimento Científico e Tecnológico (CNPq, grants 304291/2020-0 and 408764/2024-4), and the Fundação de Amparo à Pesquisa do Estado de São Paulo (FAPESP, grant 2021/03868-8). L.T.S. thanks FAPESP (14/23504-7, 17/12716-1) and Faepex (convênio 519.292 solicitação 059/15) fellowships, and a Center for Systematic Entomology grant in 2018. This project was partially funded through grants from the University of Florida’s Biodiversity Institute, the Research Opportunity Seed Fund (#00133613), and the SEED AI award (#00133688) to AP. We are also grateful to Akito Kawahara and Joe Martinez for initial discussions on butterfly biology and potential approaches. Finally, we thank the Research Software Engineering and the NVIDIA AI Technology Center (NVAITC) teams at UF for their support of this project.

## Data availability statement

We deposited the images (segmented, downsized), the embeddings and hand-coded features, and derived traits (i.e., aposematic scores and similarity metrics), as well as all code and scripts required to reproduce the figures and the results in an online repository: https://doi.org/10.5281/zenodo.17214905.

---

**Figure S1.**
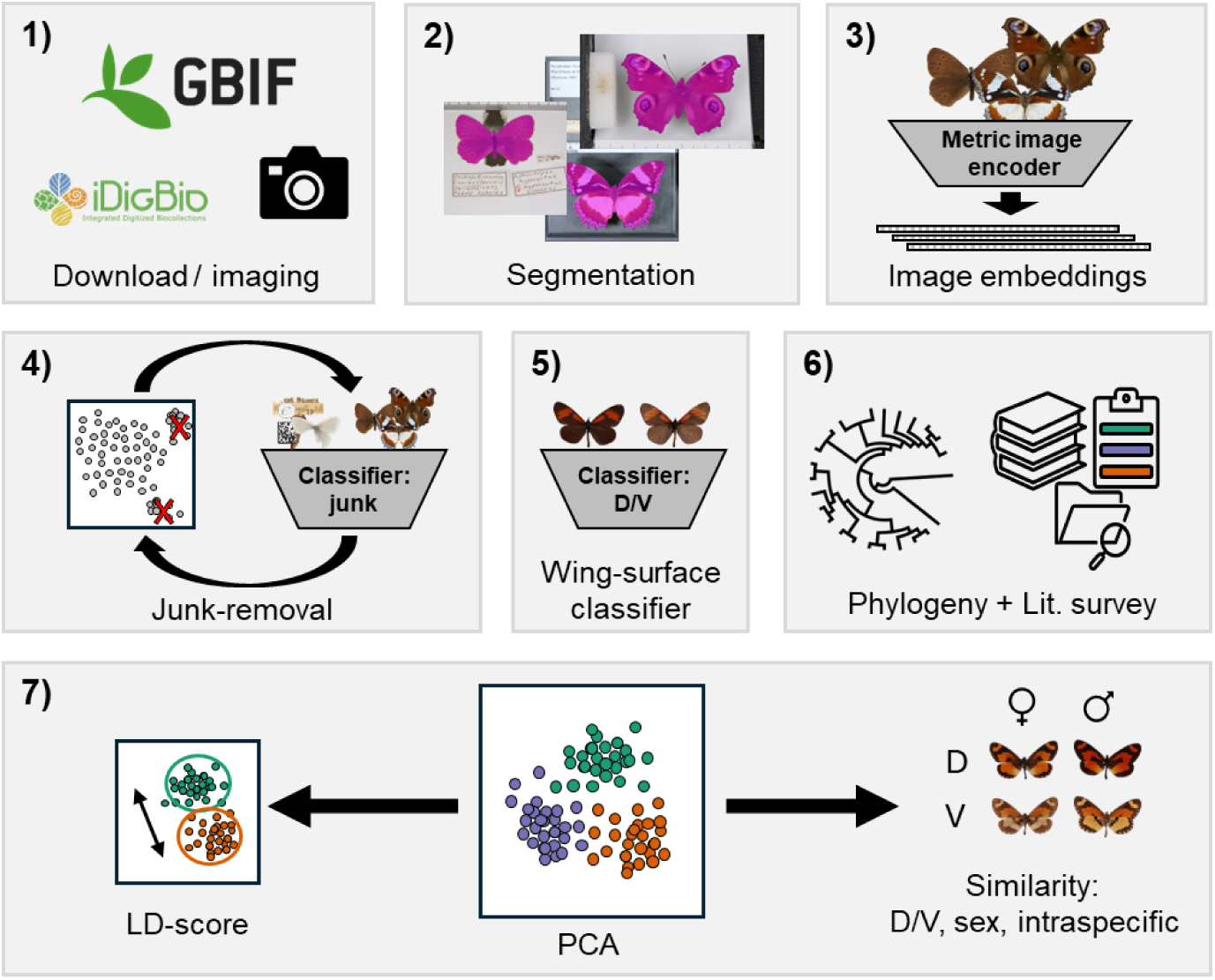
Overview of our data-collection and analytical pipeline. We assembled a large image dataset of Nymphalid butterflies by downloading 359,804 unique images from GBIF and iDigBio, and by supplementing them with 5,374 targeted photographs of 399 species (8) from major museum collections (1). We segmented specimens from all images using GroundedSAM, a foundation segmentation model (2), and converted them into 768-dimensional embedding vectors with UNICOM (30), a metric image encoder pre-trained on 400 million images (3). We visualized the embedding space with tSNE, selected unwanted junk data-points (e.g., paper labels, rulers, partial segmentations, etc.) and used them to train a YOLOv8 (80) binary classifier (99.7% accuracy) to iteratively remove all erroneous segmentations (4). A second YOLOv8 model (99.6% accuracy) assigned dorsal and ventral perspectives (5). We then combined the final, cleaned dataset of 33,450 dorso-ventral image pairs (16,725 specimens, 2,543 species) with information on chemical defense (low, medium, high) for 355 species that we collected through a structured literature survey, as well as a time-calibrated phylogeny (9) (6). Using principal component analysis (PCA), we reduced the 768-dimensional feature space to capture the main axes of wing pattern variation, yielding a comprehensive, family-wide morphospace of wing color patterns that served as the basis for all downstream analysis. Using the first seven PCs, we trained a linear discriminant model, using only the labels for “low” and “high” chemical defense and validated on an external dataset of moths, to define a univariate aposematic score (LD1). Moreover, we used both PC 1-7 (aposematic aspects only) and PC 1-177 (all phenotypic aspects) to quantify similarity across dorsal and ventral surfaces, sexes, and within species (7).

**Figure S2.**
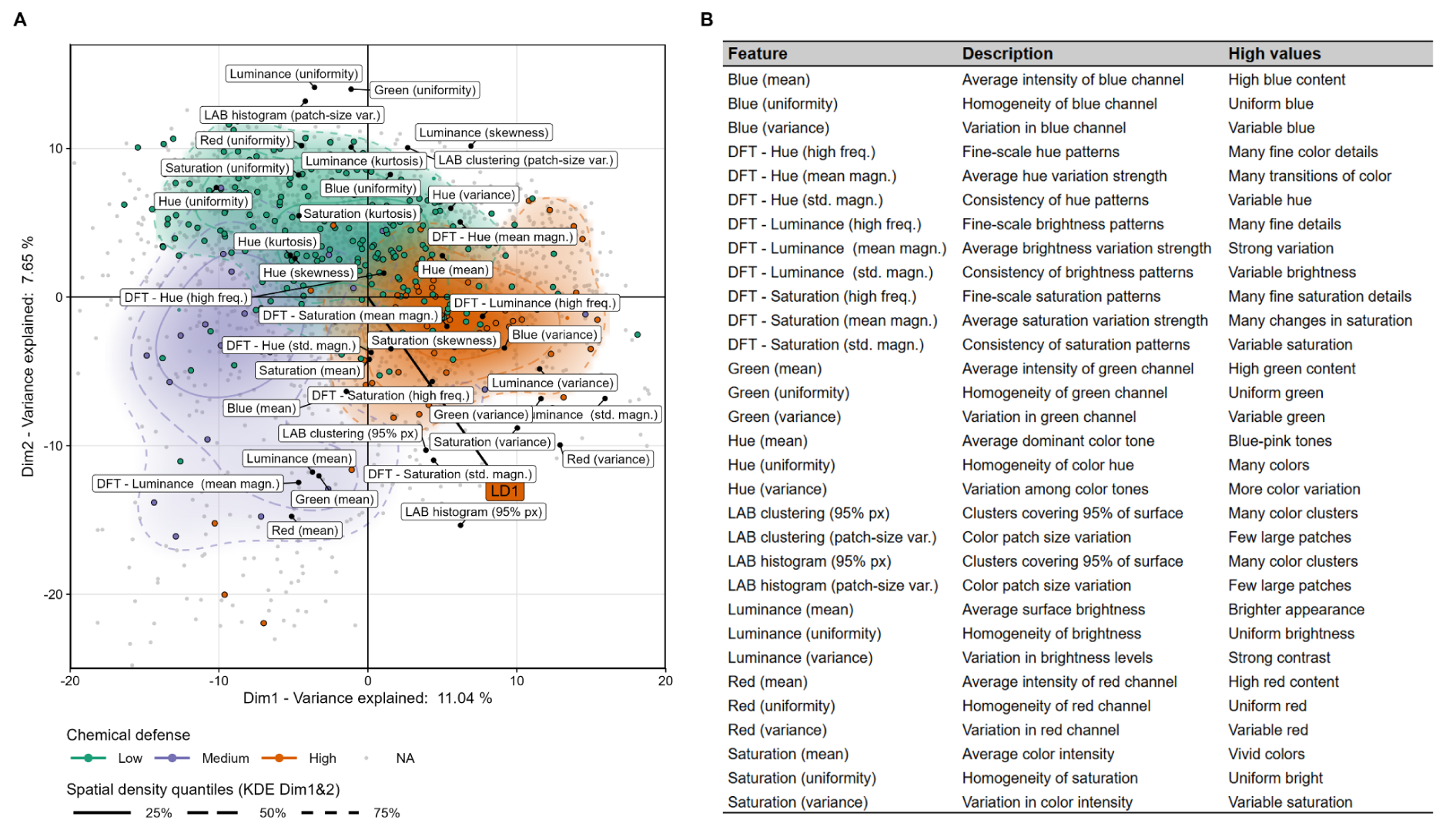
Butterfly morphospace with supplementary color pattern features A: Principal Component Analysis (PCA) of embedding vectors with supplementary color pattern features projected to illustrate associations with regions of morphospace. Points are species means across all wing surfaces and sexes are colored by chemical defense category, and contour lines show spatial density quantiles (25%, 50%, 75%). **B:** Description of color pattern features with interpretation of values (i.e., the loadings in panel A).

**Figure S3.**
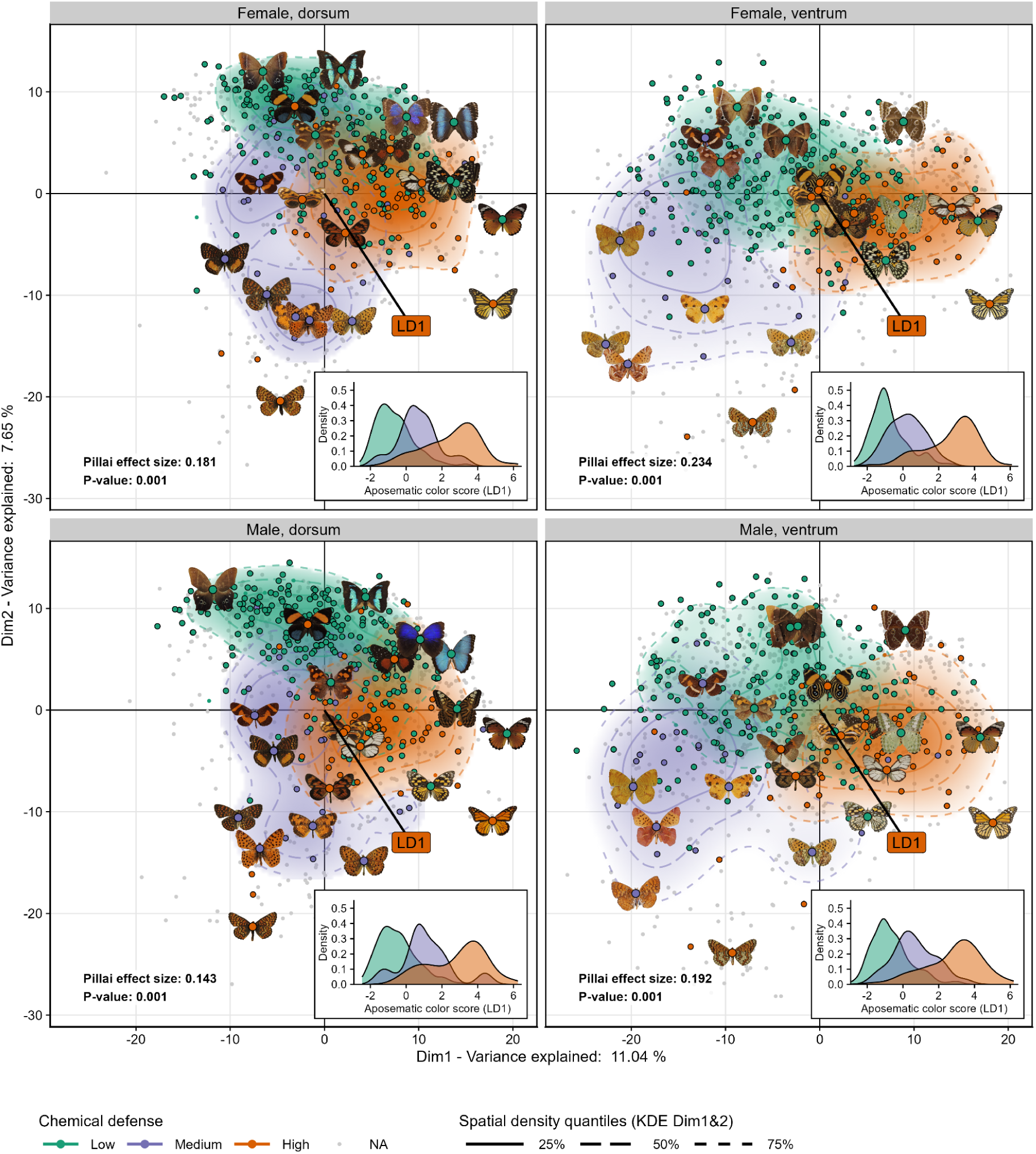
Morphospace of wing color patterns in Nymphalidae; by wing-surface and sex. Morphospace of wing color patterns constructed from the same Principal Component Analysis (PCA) as shown in Fig. 1, but aggregated to centroids for wing surface and sex for each species (N=1095, Table 1). Species categorized as low, medium, or high chemical defense separate in multivariate space, as confirmed by a phylogenetically informed multivariate GLS (Table S1A; P<0.001 for all wing surface and sex contexts). Strongest separation occurs in females on the ventrum (23.4% variation explained), and weakest separation for males on the dorsum (14.3% explained).

**Figure S4.**
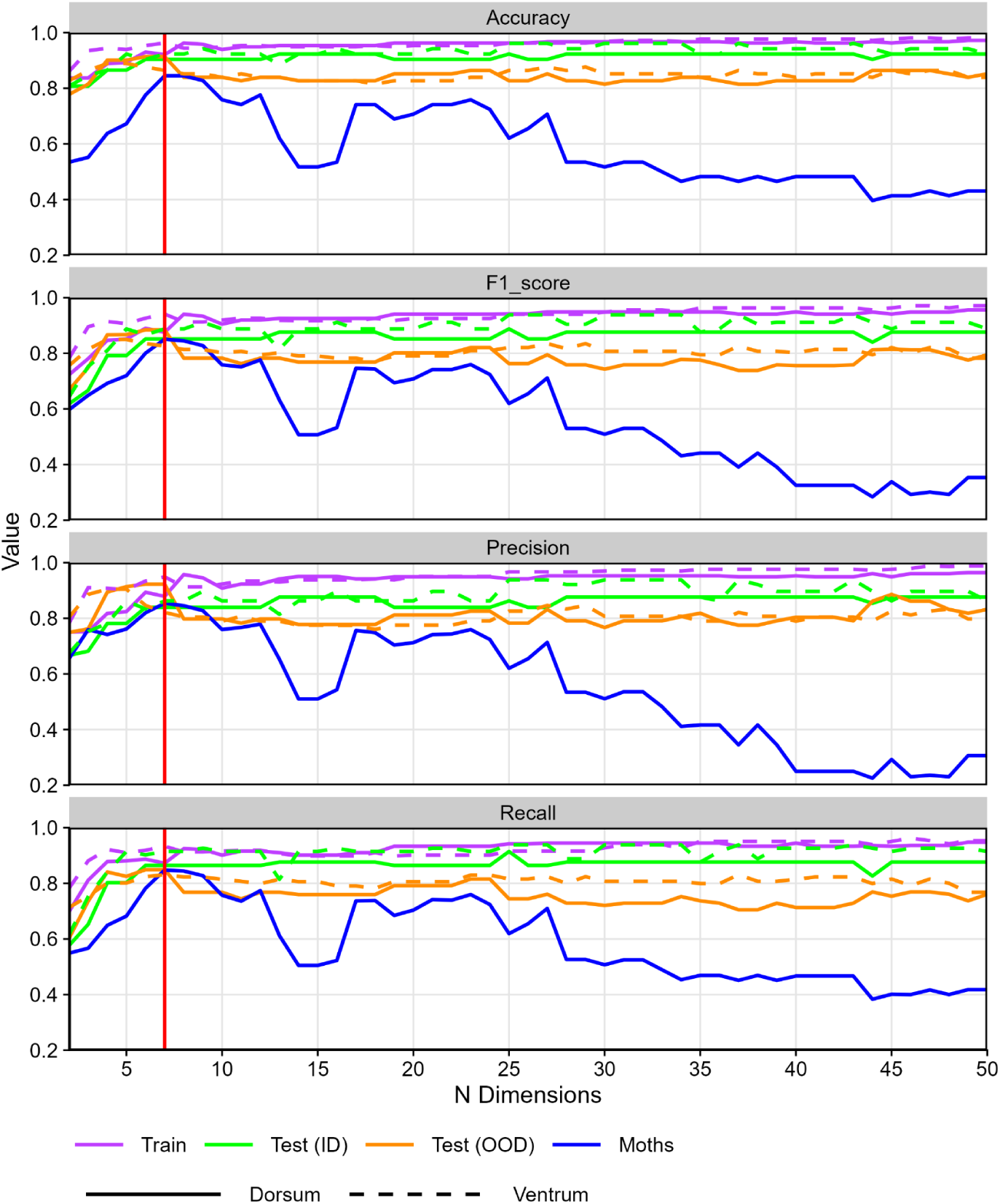
Performance of linear discriminant analysis (LDA) Results for an LDA classifier trained on different numbers of principal components (PCs) for the dorsal side across different datasets. Shown are accuracy, F1-score, precision, and recall for the training set (purple), in-distribution test species from the same pool (ID, green), out-of-distribution butterfly species excluded prior to creating the train/test split (OOD, orange), and an external moth dataset (blue). Solid and dashed lines correspond to dorsum and ventrum surfaces, respectively (moths are dorsal only). The red vertical line indicates the chosen final model dimensionality (seven PCs).

**Figure S5.**
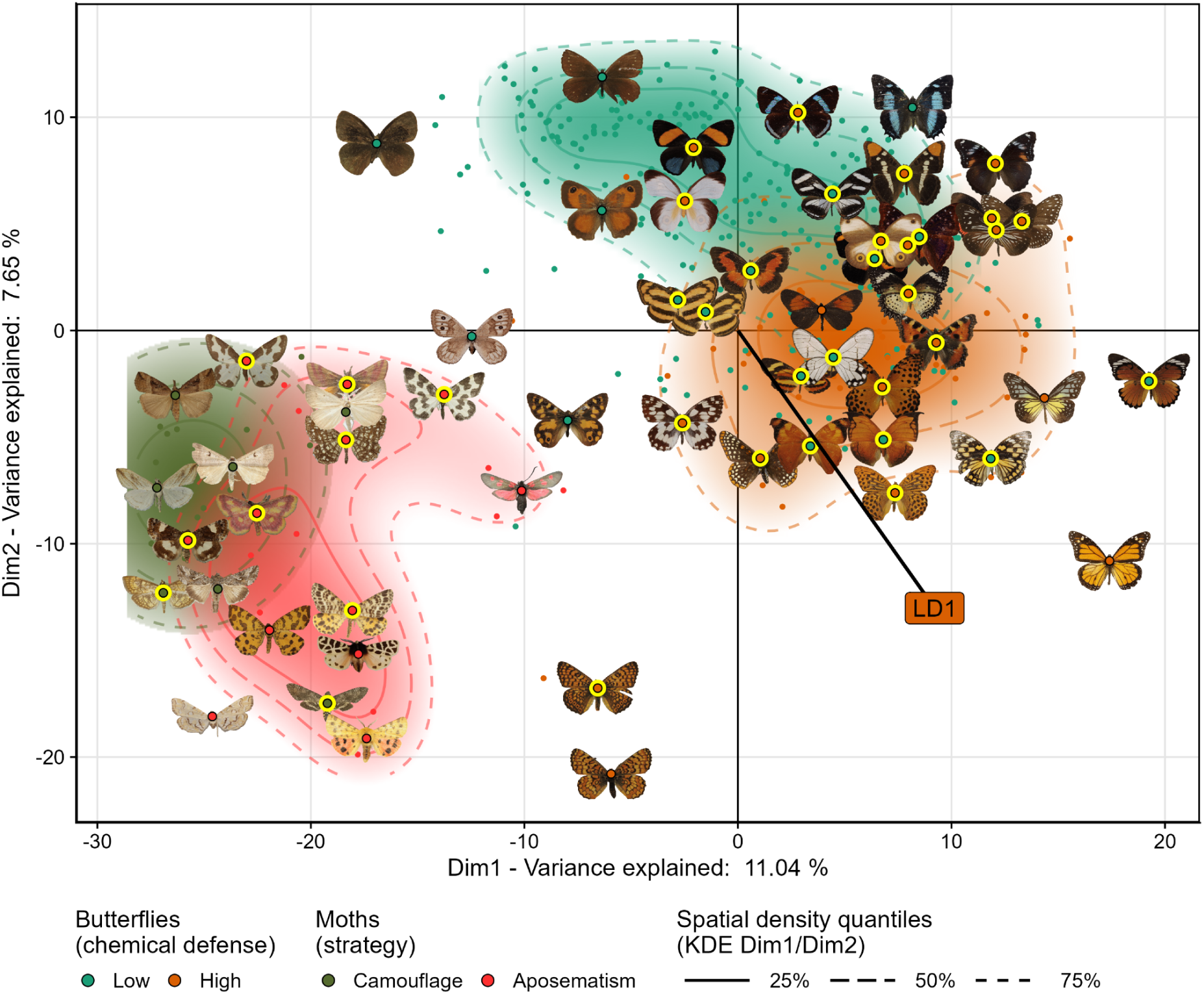
Validation of the aposematic axis using expert labelled moths. Morphospace of wing color patterns constructed from the same Principal Component Analysis (PCA) as shown in Fig. 1 (dorsal only), but together with moth species projected into the same PC space (color coded by signaling strategy). Each point represents the species centroid for the dorsum in PC space; yellow points highlight cases where class assignment did not match the expected category (see Table S5). Pictograms illustrate these cases, along with 10 representative species associated with different regions of morphospace. Contour lines denote spatial density quantiles (25%, 50%, 75%). The discriminant axis (LD1) aligns with the separation between low- and high-defense butterflies, and aposematic versus camouflaged moths segregate along the same axis in morphospace, supporting the interpretation of LD1 as an axis of aposematic signaling.

**Figure S6.**
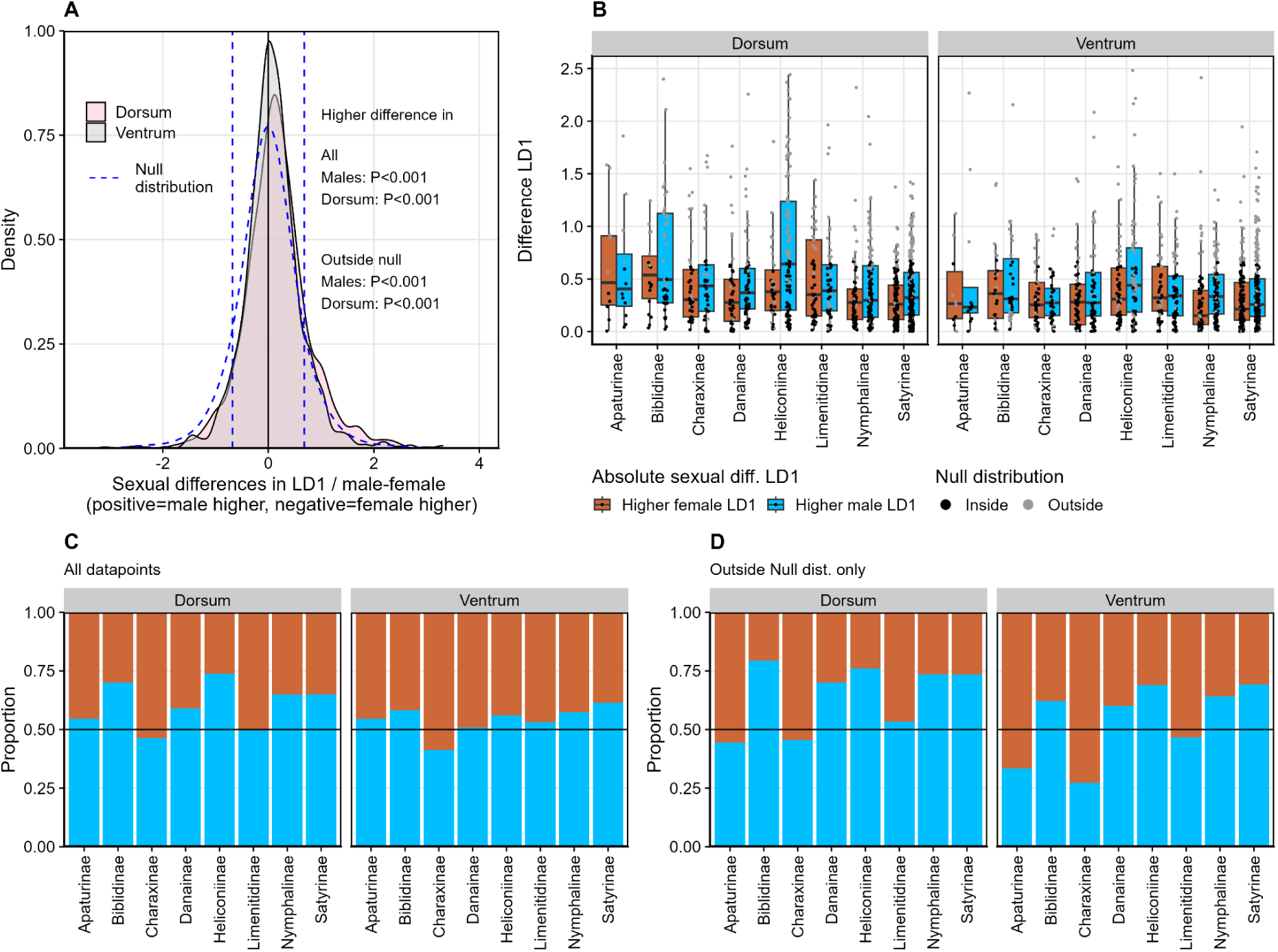
Sexual differences in aposematic score A: Distribution of sexual differences in aposematic score (LD1) for dorsal and ventral side, as well as the expected null distribution (1000 simulations of random chance for N=2190 observations (D/V)) and its 10/90% cutoff. Positive values indicate higher male score, negative higher female score. Shown P-values are the results of two phylogenetic regressions (Table S4), which indicate higher differences in LD1 for males and on the dorsal wing surface (main effect only), both for all species and those outside the null distribution. **B:** Boxplots show absolute differences in LD1 for males and females, broken down by subfamily and wing surface. A species was assigned to be outside the null distribution if as much as one wing surface was outside the cutoff (panel A). **C:** Proportion of species per family with higher male or female LD1, indicating a general male bias toward higher aposematic scores (P<0.001; Table S3). **D:** Proportion of species only outside the null distribution. In those species with higher than expected differences in LD1, the male bias was even stronger (P<0.001; Table S3).

**Figure S7.**
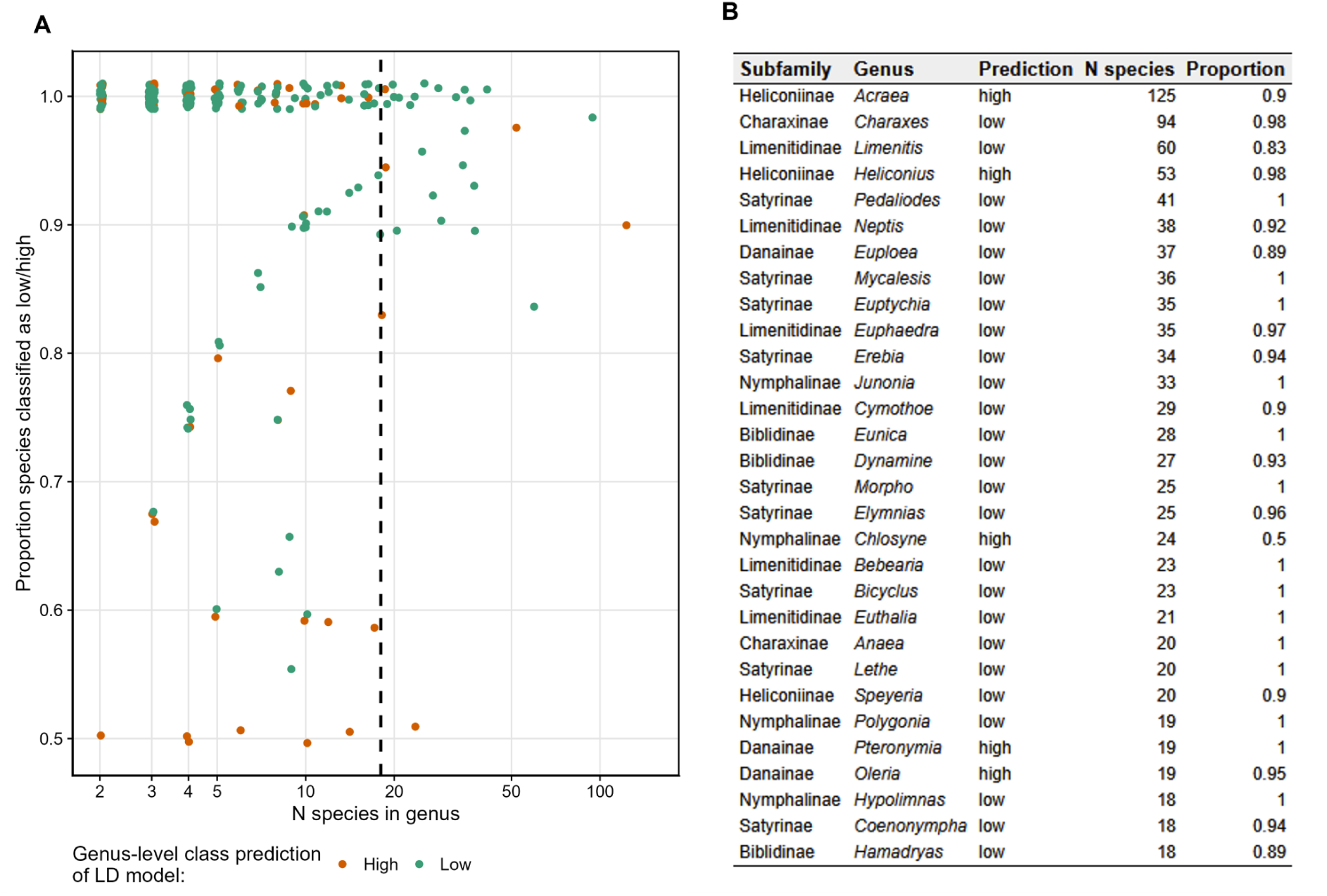
Genus-level consistency in the assignment of aposematic scores A: The proportion of species-level class assignments per genus as a function of genus size (dashed line = 18, genus size cutoff for Table in panel B) across the entire dataset, i.e., all genera (labeled and unlabeled) represented with at least two species (N=288). **B:** The 30 largest genera. The LD-model was highly consistent at the genus level: 227 out of 288 genera (81%) were perfectly consistent in their class assignment, 241 genera (86%) had >90% of species assigned to the same class, and 279 genera (97%) had >80% congruence.

**Figure S8.**
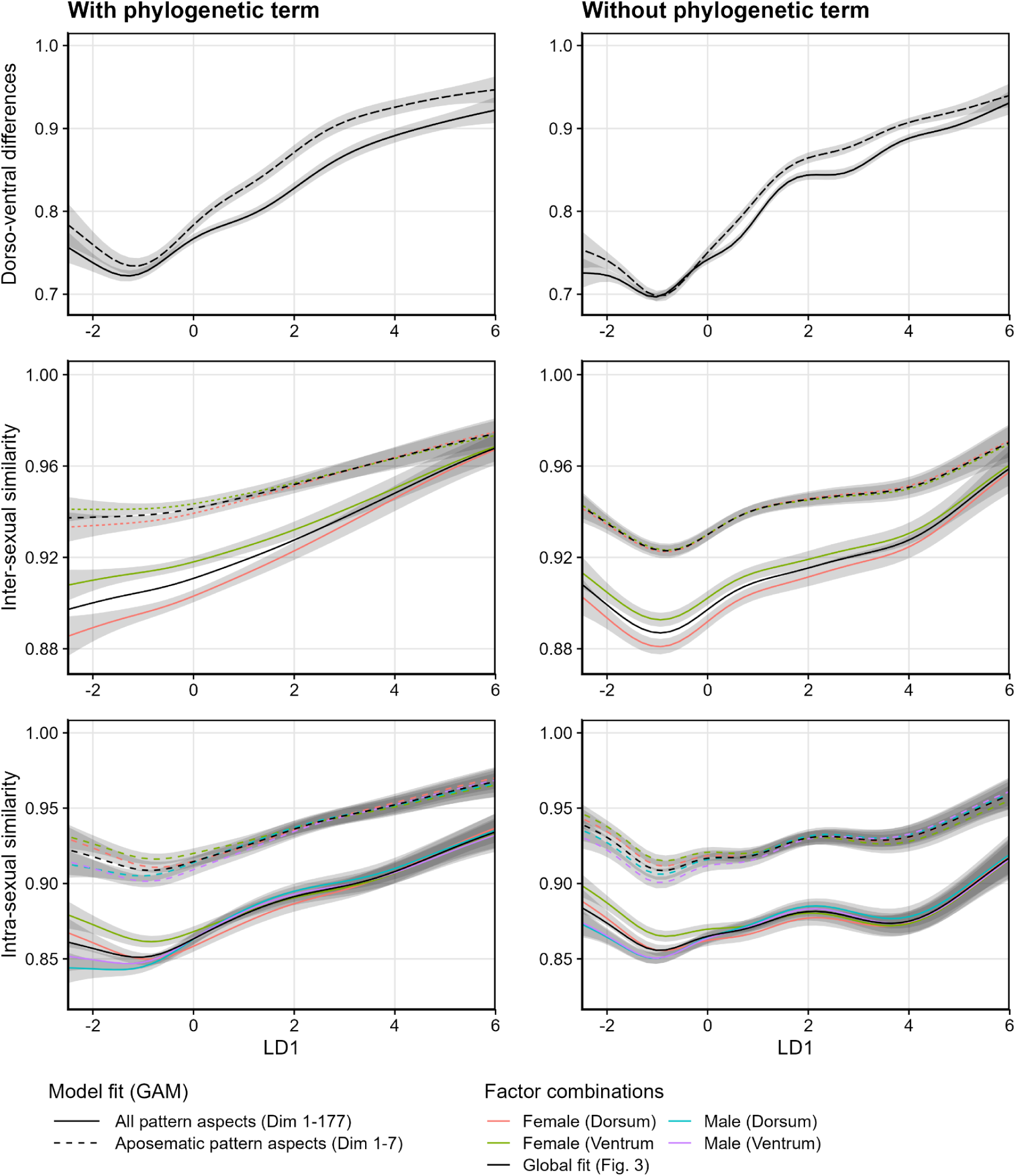
Partial model fits for similarity regressions. Like in Fig. 3, each row/panel shows the relationship between aposematic score (LD1) and different axes of intraspecific phenotypic similarity. Solid lines represent fits from Generalized Additive Models (GAM) based on all pattern aspects (PCs 1–177; Table 2), while dashed lines represent GAM fits based only on aposematic aspects (PCs 1–7). The left column corresponds to the data presented in Fig. 3, i.e., GAMs with a phylogenetic error term, whereas the right column includes more species that, however, were not present in the phylogenetic tree (Table S4; global [black] and partial fits [color coded]).

**Figure S9.**
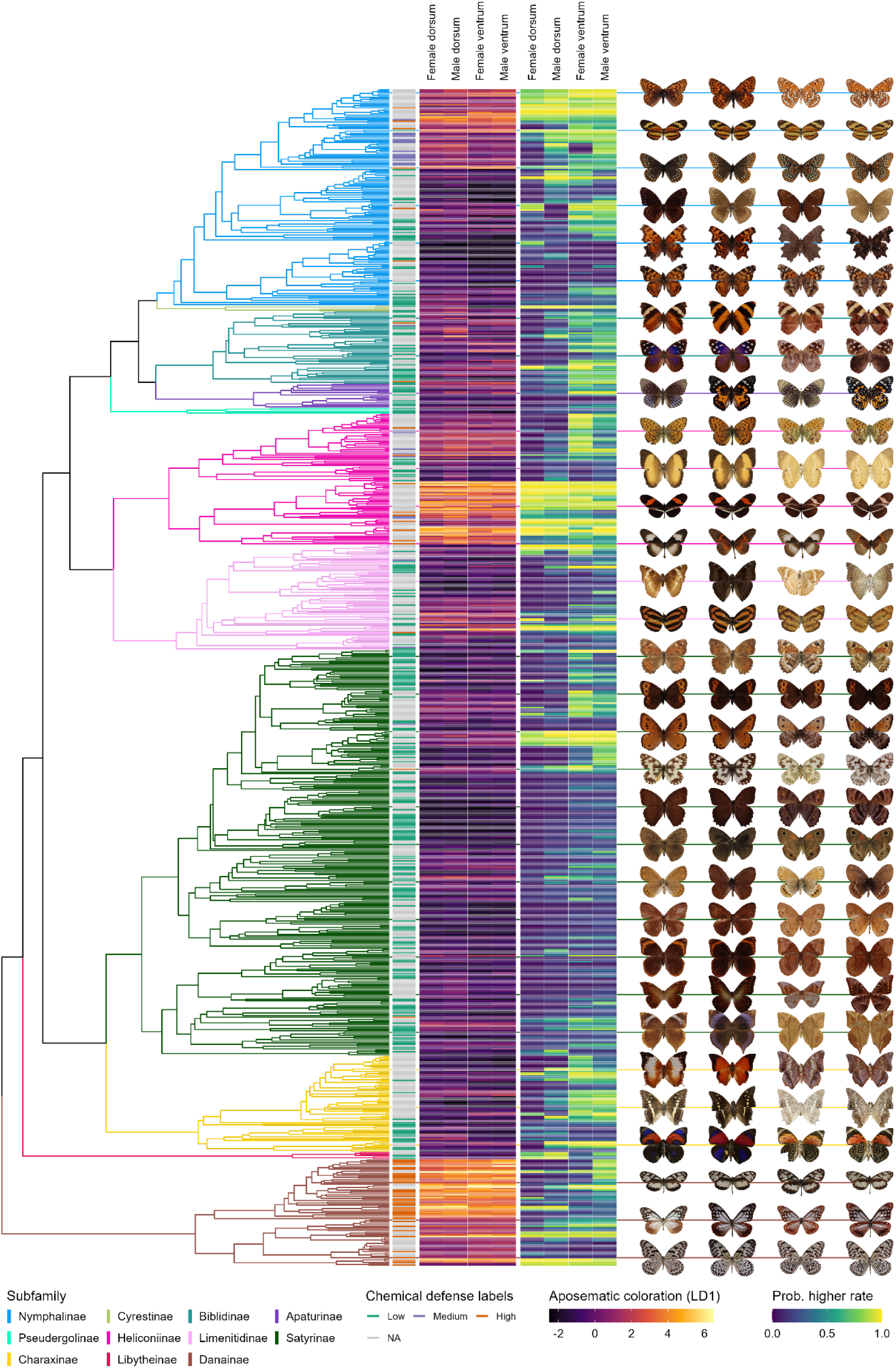
Aposematic scores and evolutionary rates by sex and wing surface. Expanding Fig. 4, shown are - from left to right - the phylogenetic tree (color coded by subfamily) for all species with sex-information in the phylogeny (N=783; including species without labels for chemical defense), aposematic score for each wing-surface- and sex-context, the estimated probabilities for a higher evolutionary rate, and selected examples (same order as the score and rate columns).

**Figure S10.**
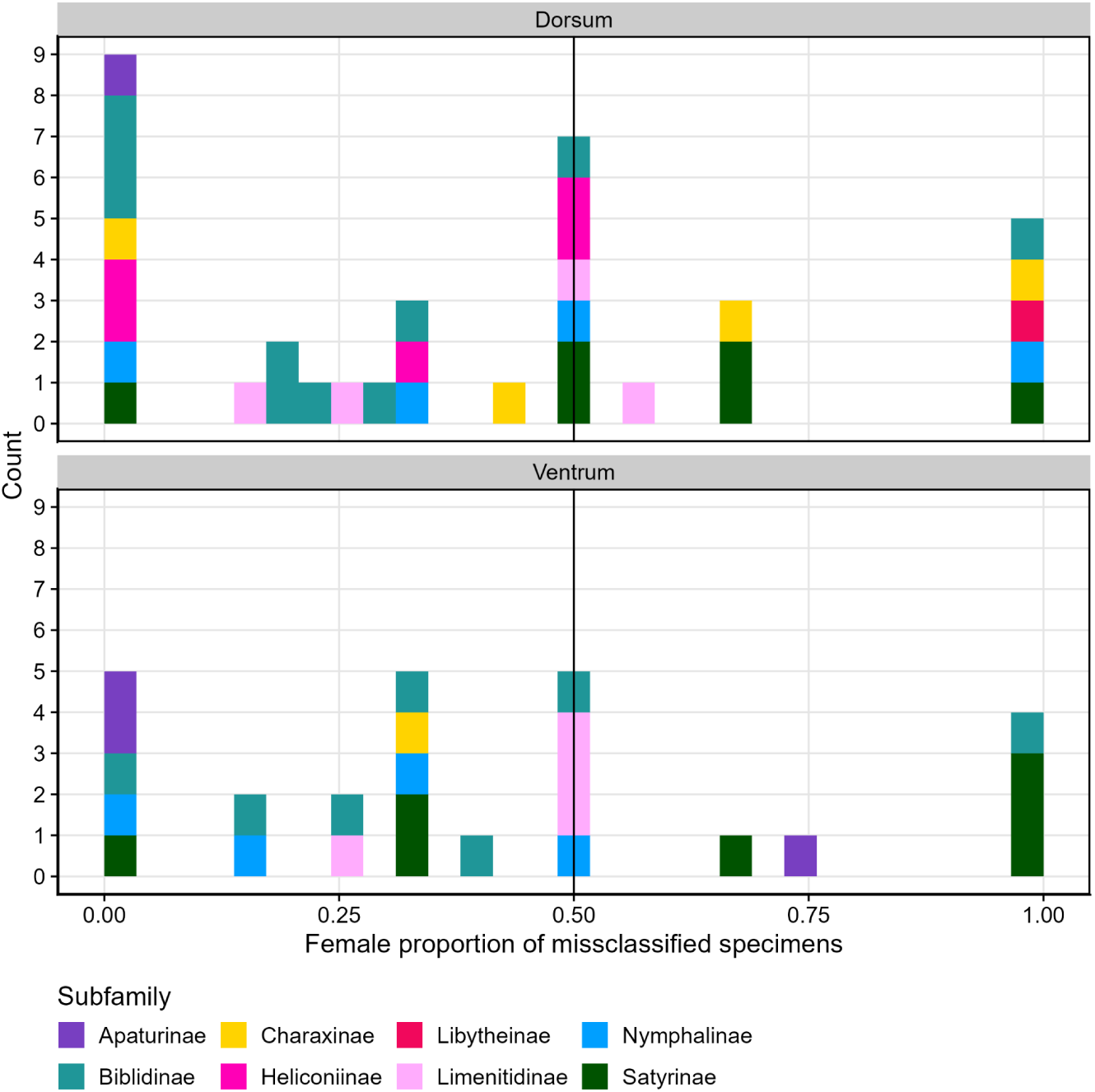
Sex bias in misclassified specimens across Nymphalidae subfamilies. Histograms showing the female proportion of misclassified specimens for each species on the dorsum (top) and ventrum (bottom), with bars colored by subfamily (also see Table S6). A vertical line at 0.5 indicates equal proportions of female and male misclassifications. Most cases show no consistent female bias, with scattered instances across subfamilies and wing surfaces. Note that this is across all misclassified specimens, not just those belonging to species that were misclassified as a whole (Table S5).

**Table S1.** Results of multivariate phylogenetic analysis A: Results from multivariate phylogenetic generalized least squares (mvGLS) models testing the separation by chemical defense in multivariate space. The models used PCs 1-177, aggregated to the species centroids for each species across all wing surfaces and sexes (mvGLS; shown in Fig. S2), aggregated to species-centroids for dorsum and ventrum (mvGLS 2; shown in Fig. 1), and for sex by wing surface (mvGLS 3; shown in Fig. S3). Reported are Pillai’s trace statistics, permutation-based P-values, and effect sizes. All models show highly significant phenotypic differences among groups in embedding space. **B:** Estimating the strength of the phylogenetic signal using Bloomberg’s k, for labeled species (same N as for A), and for all species in the phylogeny (ext=extended).

| **A** |  |  |  |  |  |  | **B** |  |  |  |  |
| --- | --- | --- | --- | --- | --- | --- | --- | --- | --- | --- | --- |
| Model name | Wing surface | Sex | N | Pillai statistic | Effect size | P-value | Bloomberg's k LD1 | P-value | N (ext.) | Bloomberg's k LD1 (ext) | P-value (ext) |
| mvGLS 1 | - | - | 355 | 0.887 | 0.187 | <0.001 | 1.132 | 0.001 | 1382 | 0.593 | 0.001 |
| mvGLS 2a | dorsum | - | 355 | 0.830 | 0.144 | <0.001 | 1.160 | 0.001 | 1382 | 0.461 | 0.001 |
| mvGLS 2b | ventrum | - |  | 0.917 | 0.207 | <0.001 | 0.904 | 0.001 |  | 0.539 | 0.001 |
| mvGLS 3a | dorsum | female | 336 | 0.902 | 0.181 | <0.001 | 1.047 | 0.001 | 783 | 0.488 | 0.001 |
| mvGLS 3b | dorsum | male |  | 0.854 | 0.143 | <0.001 | 1.094 | 0.001 |  | 0.597 | 0.001 |
| mvGLS 3c | ventrum | female |  | 0.975 | 0.234 | <0.001 | 0.882 | 0.001 |  | 0.626 | 0.001 |
| mvGLS 3d | ventrum | male |  | 0.915 | 0.192 | <0.001 | 0.864 | 0.001 |  | 0.683 | 0.001 |

**Table S2.**
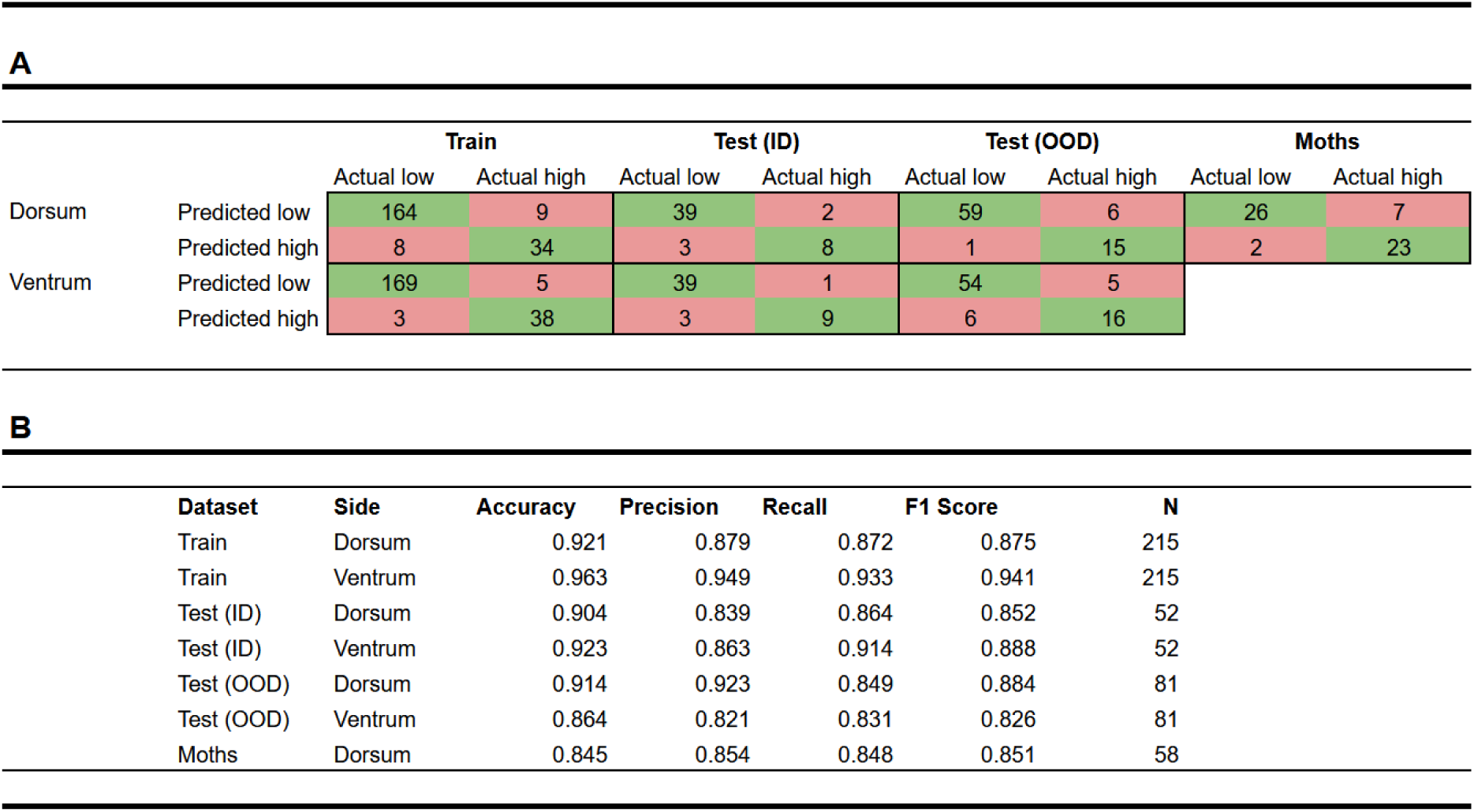
Confusion matrix and performance metrics for linear discriminant classifier A: Confusion matrices showing classification performance of the final linear discriminant (LD) model across datasets (from left to right): training, in-distribution test (ID), out-of-distribution test (OOD), and external moth dataset, separated by wing surface (dorsum, ventrum). Values indicate the number of specimens predicted as high vs. low aposematic class relative to their actual class, showing that the final classifier was robust across contexts. **B:** Summary statistics for each dataset including accuracy, precision, recall, F1 score, and sample size (N). The final model with seven components (for all tested models see Fig. S3) achieved high accuracy and balanced performance across datasets, with slightly lower but still robust performance when applied to moths.

|  |  | Train |  | Test (ID) |  | Test (OOD) |  | Moths |  |
| --- | --- | --- | --- | --- | --- | --- | --- | --- | --- |
|  |  | Actual low | Actual high | Actual low | Actual high | Actual low | Actual high | Actual low | Actual high |
| Dorsum | Predicted low | 164 | 9 | 39 | 2 | 59 | 6 | 26 | 7 |
|  | Predicted high | 8 | 34 | 3 | 8 | 1 | 15 | 2 | 23 |
| Ventrums | Predicted low | 169 | 5 | 39 | 1 | 54 | 5 |  |  |
|  | Predicted high | 3 | 38 | 3 | 9 | 6 | 16 |  |  |

**B**
| Dataset | Side | Accuracy | Precision | Recall | F1 Score | N |
| --- | --- | --- | --- | --- | --- | --- |
| Train | Dorsum | 0.921 | 0.879 | 0.872 | 0.875 | 215 |
| Train | Ventrums | 0.963 | 0.949 | 0.933 | 0.941 | 215 |
| Test (ID) | Dorsum | 0.904 | 0.839 | 0.864 | 0.852 | 52 |
| Test (ID) | Ventrums | 0.923 | 0.863 | 0.914 | 0.888 | 52 |
| Test (OOD) | Dorsum | 0.914 | 0.923 | 0.849 | 0.884 | 81 |
| Test (OOD) | Ventrums | 0.864 | 0.821 | 0.831 | 0.826 | 81 |
| Moths | Dorsum | 0.845 | 0.854 | 0.848 | 0.851 | 58 |

**Table S3.**
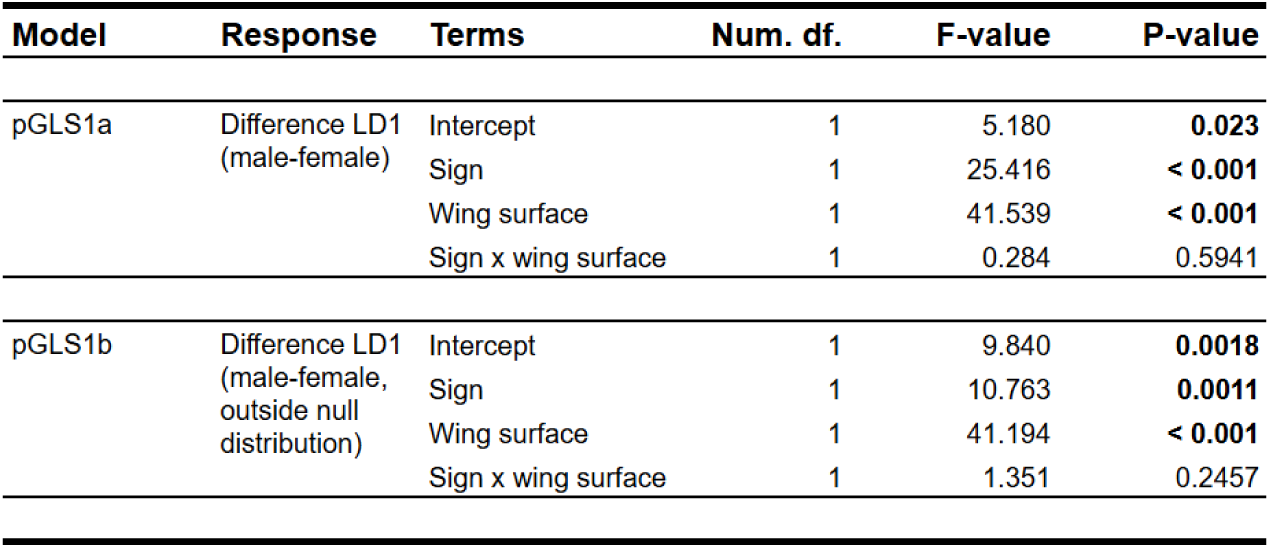
Results for phylo-regression of intensity of sex-bias in aposematic score. Results from phylogenetic GLS models testing differences in aposematic scores (LD1) between males and females across wing surfaces (Fig. S7). Models include the sign of the sex difference (male higher vs. female higher), wing surface (dorsal vs. ventral), and their interaction. We fit one model on the full dataset; (pGLS1a) and one model restricted to species with sex differences outside the null distribution (pGLS1b). Significant effects of both sign and wing surface indicate systematic sex-related variation in LD1 scores, while the interaction term was not significant.

| **Model** | **Response** | **Terms** | **Num. df.** | **F-value** | **P-value** |
| --- | --- | --- | --- | --- | --- |
| pGLS1a | Difference LD1<br>(male-female) | Intercept | 1 | 5.180 | **0.023** |
|  |  | Sign | 1 | 25.416 | **< 0.001** |
|  |  | Wing surface | 1 | 41.539 | **< 0.001** |
|  |  | Sign x wing surface | 1 | 0.284 | 0.5941 |
| pGLS1b | Difference LD1<br>(male-female,<br>outside null<br>distribution) | Intercept | 1 | 9.840 | **0.0018** |
|  |  | Sign | 1 | 10.763 | **0.0011** |
|  |  | Wing surface | 1 | 41.194 | **< 0.001** |
|  |  | Sign x wing surface | 1 | 1.351 | 0.2457 |

**Table S4.**
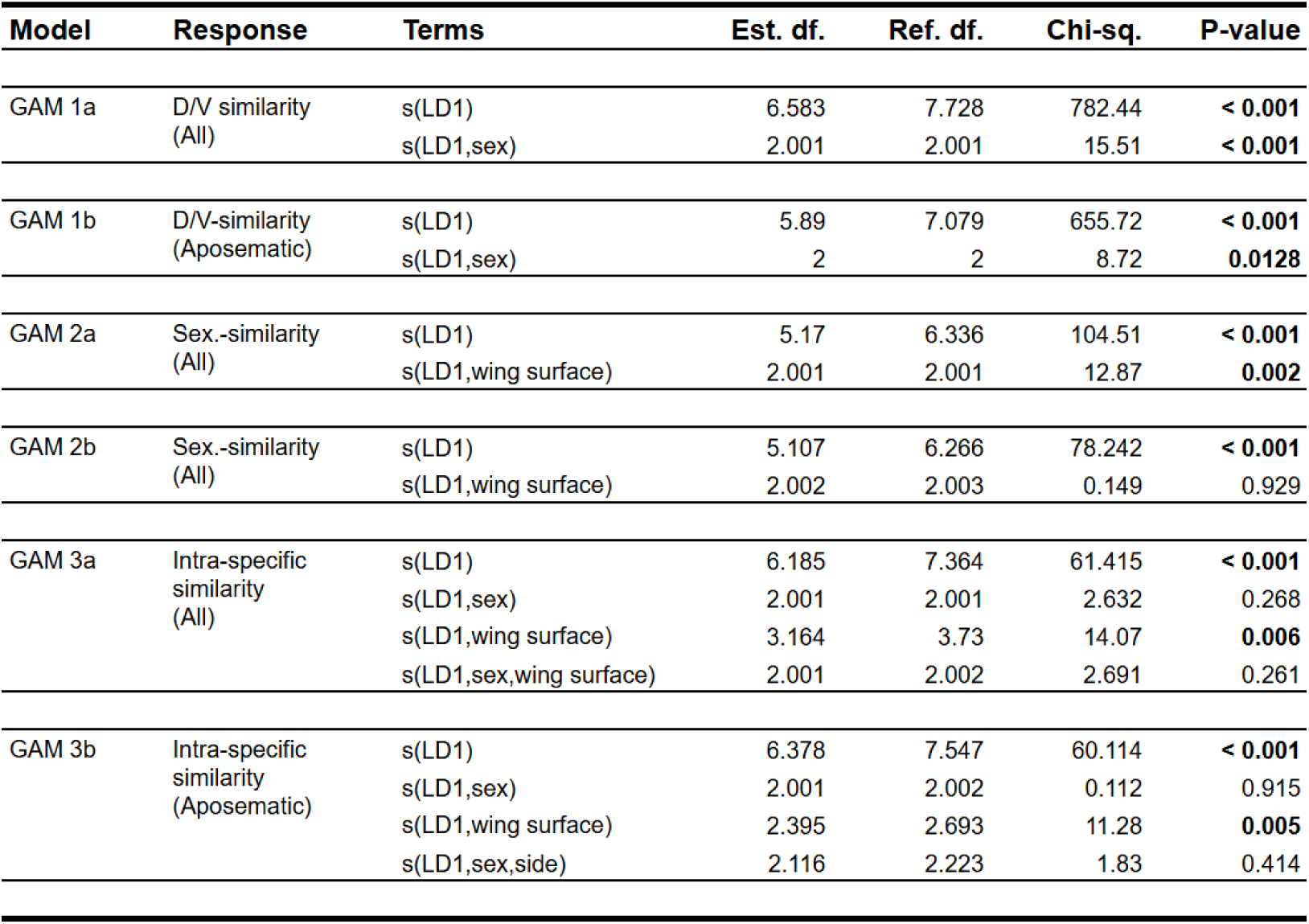
Results for aposematic score and phenotypic similarity (non-phylo. models) Results for Generalized Additive Models (GAM) of phenotypic similarity (dorso-ventral, sexual, and intra-specific) over aposematic score (LD1). All shown models use the betar family (logit function) and 10 knots per term. Shown columns are, from left to right: model names, response variables (All=similarity based on PC1-177, Aposematic=similarity based on PC1-7), terms for main-effect- (s(LD1)) and factor-interaction-smooths (s(LD1), …), estimated degrees of freedom (Est. df.), reference degrees of freedom Ref. df.), Chi-squared statistic (Chi-sq.), and P-value (significant >=0.05 in bold). These models are the non-phylogenetic counterparts with higher sample sizes (Table 1, Fig. S6) to the models shown in the main text (Fig. 3, Table 2).

**Table S5.** Species with divergent class assignments. Listed are species that counted as misclassified, i.e., when the linear discriminant model assigned the species-centroid to a different class than our label for chemical defense. The columns show (from left to right) the total number of specimens vs. misclassified specimens, and further separation by wing surface and sex (a specimen counted as misclassified when at least one surface was divergent). Overall, classification accuracy was high (training set: 92% on the dorsum and 96% on the ventrum, Table S2). Some of these cases are well-known examples of Batesian mimicry, such as *Pseudacraea poggei* (False Monarch) and *Eresia eunice* (Tiger Crescent). In other species, including *Lasippa tiga* and *Catonephele numilia*, divergence occurred only on one surface (in these cases, the dorsum). For more detailed information on intra-specific misclassifications, with a focus on female-limited Batesian mimicry of aposematic color patterns, see Table S6.

| Subfamily | Species | Specimens<br>total/misscl. | Dorsum<br>f/m/u | Ventrum<br>f/m/u |
| --- | --- | --- | --- | --- |
| Low chemical defense - classified as "high" |  |  |  |  |
| Apaturinae | *Hestina assimilis* | 2/1 | 0/1/1 | 0/0/0 |
| Apaturinae | *Sephisia dichroa* | 10/6 | 2/7/1 | 0/0/0 |
| Biblidinae | *Batesia hypochlora* | 10/10 | 0/0/0 | 4/4/2 |
| Biblidinae | *Catonephele numilia* | 17/14 | 6/10/1 | 0/0/0 |
| Biblidinae | *Mestra hypermestra* | 10/6 | 0/0/0 | 4/5/1 |
| Biblidinae | *Temenis laothoe* | 14/10 | 3/6/5 | 0/0/0 |
| Biblidinae | *Vila azeka* | 8/8 | 2/6/0 | 2/6/0 |
| Charaxinae | *Coenophlebia archidona* | 11/9 | 3/5/3 | 0/0/0 |
| Limenitidinae | *Crenidomimas concordia* | 11/6 | 0/0/0 | 3/8/0 |
| Limenitidinae | *Lasippa tiga* | 34/34 | 4/3/27 | 0/0/0 |
| Limenitidinae | *Neptis ida* | 9/8 | 0/0/0 | 5/4/0 |
| Limenitidinae | *Pantoporia sandaka* | 17/16 | 3/3/11 | 0/0/0 |
| Limenitidinae | *Pseudacraea poggei* | 8/8 | 2/6/0 | 2/6/0 |
| Nymphalinae | *Eresia eunice* | 8/8 | 3/3/2 | 3/3/2 |
| Nymphalinae | *Vanessula milca* | 15/12 | 6/9/0 | 0/0/0 |
| Satyrinae | *Argyrophenga antipodium* | 7/6 | 0/0/0 | 3/4/0 |
| Satyrinae | *Cithaerias aurora* | 1/1 | 0/0/0 | 0/1/0 |
| Satyrinae | *Erebiola butleri* | 5/3 | 0/0/0 | 3/2/0 |
| Satyrinae | *Pseudohaetera hypaesia* | 5/5 | 2/2/1 | 2/2/1 |
| High chemical defense - classified as "low" |  |  |  |  |
| Biblidinae | *Biblis hyperia* | 4/4 | 2/2/0 | 2/2/0 |
| Biblidinae | *Callicore tolima* | 14/12 | 3/8/3 | 0/0/0 |
| Biblidinae | *Diaethria clymena* | 16/16 | 5/9/2 | 5/9/2 |
| Danainae | *Anetia briarea* | 7/7 | 3/4/0 | 3/4/0 |
| Danainae | *Euploea camaralzeman* | 20/16 | 3/8/9 | 0/0/0 |
| Danainae | *Parantica luzonensis* | 10/10 | 4/4/2 | 0/0/0 |
| Heliconiinae | *Argynnis paphia* | 15/15 | 5/8/2 | 5/8/2 |
| Heliconiinae | *Cethosia cyane* | 8/8 | 3/5/0 | 3/5/0 |
| Limenitidinae | *Adelpha californica* | 6/6 | 3/3/0 | 3/3/0 |
| Nymphalinae | *Aglais urticae* | 10/10 | 5/5/0 | 5/5/0 |
| Nymphalinae | *Chlosyne janais* | 15/14 | 5/7/3 | 5/7/3 |
| Nymphalinae | *Euphydryas phaeton* | 117/72 | 7/14/96 | 0/0/0 |
| Nymphalinae | *Hypolimnas bolina* | 49/49 | 14/8/27 | 14/8/27 |
| Nymphalinae | *Melitaea cinxia* | 13/7 | 6/6/1 | 0/0/0 |
| Satyrinae | *Hyantis hodeva* | 5/5 | 2/3/0 | 2/3/0 |
| Satyrinae | *Melanargia galathea* | 17/14 | 5/12/0 | 5/12/0 |
| Satyrinae | *Taenaris catops* | 4/4 | 2/2/0 | 2/2/0 |

**Table S6.**
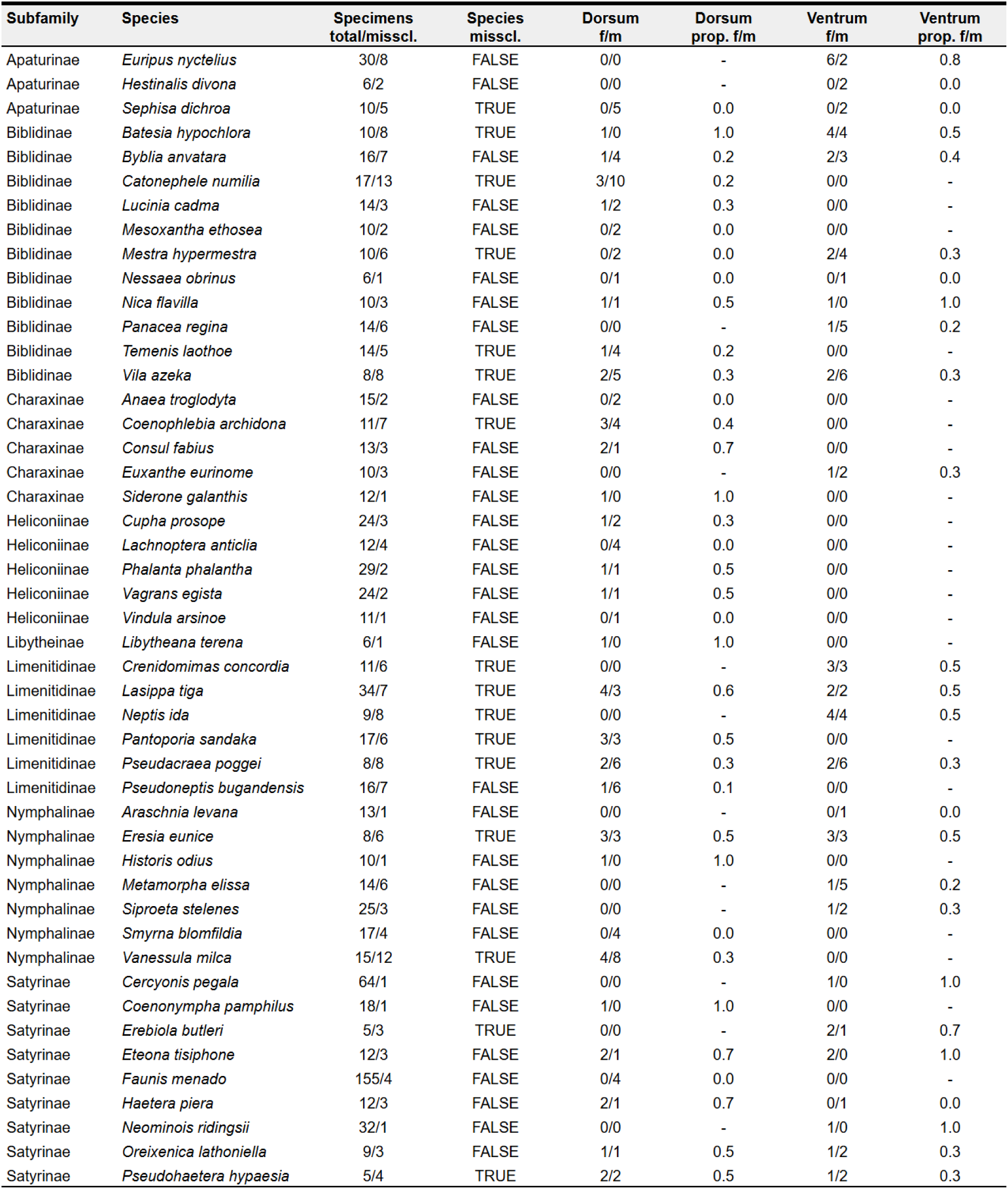
Species with divergent class assignments (low chemical defense & complete sex-information only) Listed are species labeled as having low chemical defense that contained any specimens with divergent class assignments in the linear discriminant model, focussing only those specimens with complete sex information (i.e., no “unkown”). The columns show (from left to right) the total number of specimens vs. the number of misclassified specimens, and whether misclassification occurred at the species level (as listed in Table S5). Misclassified specimens are further separated by wing surface (dorsum vs. ventrum) and sex, with female/male ratios and the proportion of females among misclassified individuals (corresponding to Figure S9). Some of these are well known cases of mimicry, such as *Consul fabus* (dorsal only) or *Eteona tisiphone*.

## References

1. D. J. Kemp, et al., An integrative framework for the appraisal of coloration in nature. Am. Nat. 185, 705–724 (2015).

2. H. W. Bates, Proposal of aposematic function of bright color patterns and Batesian mimicry. Contributions to an insect fauna of the Amazon valley. Lepidoptera: Heliconidae. Transactions Linnean Society London 23, 495–566 (1862).

3. A. R. Wallace, Mimicry, and Other Protective Resemblances Among Animals. Westminster Review (1867).

4. C. Darwin, The descent of man, and Selection in relation to sex, Vol 1 (John Murray, 1871).

5. J. J. Wiens, Z. Emberts, How life became colourful: colour vision, aposematism, sexual selection, flowers, and fruits. Biol. Rev. Camb. Philos. Soc. (2024). 10.1111/brv.13141.

6. Y. Sondhi, E. A. Ellis, S. M. Bybee, J. C. Theobald, A. Y. Kawahara, Light environment drives evolution of color vision genes in butterflies and moths. *Commun*. Biol. 4, 177 (2021).

7. D. Grimaldi, M. S. Engel, Cambridge evolution series: Evolution of the insects (Cambridge University Press, 2005).

8. N. Wahlberg, et al., Nymphalid butterflies diversify following near demise at the Cretaceous/Tertiary boundary. Proc. Biol. Sci. 276, 4295–4302 (2009).

9. N. Chazot, et al., Conserved ancestral tropical niche but different continental histories explain the latitudinal diversity gradient in brush-footed butterflies. Nat. Commun. 12, 5717 (2021).

10. B. N. Schwanwitsch, On the ground plan of wingpattern in Nymphalids and certain other families of the Rhopalocerous Lepidoptera. Proceedings Zoological Society London 34, 509–528 (1924).

11. F. Süffert, Zur Vergleichenden Analyse der Schmetterlingszeichnung. Biologisches Zentralblatt 47, 385 (1927).

12. H. F. Nijhout, The development and evolution of butterfly wing patterns (Smithsonian Books, 1991).

13. T. Sekimura, H. F. Nijhout, Eds., Diversity and evolution of butterfly wing patterns: An integrative approach, 1st Ed. (Springer, 2017).

14. J. F. Hoyal Cuthill, N. Guttenberg, S. Ledger, R. Crowther, B. Huertas, Deep learning on butterfly phenotypes tests evolution’s oldest mathematical model. Science Advances 5, eaaw4967 (2019).

15. J. F. Hoyal Cuthill, N. Guttenberg, B. Huertas, Male and female contributions to diversity among birdwing butterfly images. *Commun*. Biol. 7, 774 (2024).

16. A. Puissant, A. Chotard, F. L. Condamine, V. Llaurens, Convergence in sympatric swallowtail butterflies reveals ecological interactions as a key driver of worldwide trait diversification. Proc. Natl. Acad. Sci. U. S. A. 120 (2023).

17. M. P. Speed, M. A. Brockhurst, G. D. Ruxton, The dual benefits of aposematism: predator avoidance and enhanced resource collection. Evolution 64, 1622–1633 (2010).

18. L. Lindstrom, J. Mappes, R. V. Alatalo, Reactions of hand-reared and wild-caught predators toward warningly colored, gregarious, and conspicuous prey. Behav. Ecol. 10, 317–322 (1999).

19. K. L. Prudic, A. K. Skemp, D. R. Papaj, Aposematic coloration, luminance contrast, and the benefits of conspicuousness. Behav. Ecol. 18, 41–46 (2007).

20. C. G. Halpin, et al., Pattern contrast influences wariness in naïve predators towards aposematic patterns. Sci. Rep. 10, 9246 (2020).

21. L. M. Arenas, J. Troscianko, M. Stevens, Color contrast and stability as key elements for effective warning signals. Front. Ecol. Evol. 2, 86891 (2014).

22. J. L. Q. Wee, A. Monteiro, Yellow and the novel aposematic signal, red, protect *Delias* butterflies from predators. PLoS One 12, e0168243 (2017).

23. J. Mallet, M. Joron, Evolution of diversity in warning color and mimicry: Polymorphisms, shifting balance, and speciation. Annu. Rev. Ecol. Syst. 30, 201–233 (1999).

24. J. A. Endler, Signals, Signal Conditions, and the Direction of Evolution. Am. Nat. 139, S125–S153 (1992).

25. E. B. Poulton, The colours of animals: their meaning and use, especially considered in the case of insects (D. Appleton, 1890).

26. 26. G. D. Ruxton, W. L. Allen, T. N. Sherratt, M. P. Speed, “Aposematism” in Avoiding Attack: The Evolutionary Ecology of Crypsis, Aposematism, and Mimicry (2nd Edn), (Oxford University Press, 2018).

27. M. Stevens, G. D. Ruxton, Linking the evolution and form of warning coloration in nature. Proc. Biol. Sci. 279, 417–426 (2012).

28. E. S. Briolat, et al., Diversity in warning coloration: selective paradox or the norm? Biol. Rev. Camb. Philos. Soc. 94, 388–414 (2019).

29. M. D. Lürig, E. Di Martino, A. Porto, BioEncoder: A metric learning toolkit for comparative organismal biology. Ecol. Lett. 27, e14495 (2024).

30. X. An, et al., Unicom: Universal and compact representation learning for image retrieval. arXiv [cs.CV] (2023).

31. J. Clavel, G. Escarguel, G. Merceron, Mvmorph: An r package for fitting multivariate evolutionary models to morphometric data. Methods Ecol. Evol. 6, 1311–1319 (2015).

32. S. D. Finkbeiner, A. D. Briscoe, R. D. Reed, Warning signals are seductive: relative contributions of color and pattern to predator avoidance and mate attraction in *Heliconius* butterflies. Evolution 68, 3410–3420 (2014).

33. O. Nokelainen, S. A. Silvasti, S. Y. Strauss, N. Wahlberg, J. Mappes, Predator selection on phenotypic variability of cryptic and aposematic moths. Nat. Commun. 15, 1678 (2024).

34. M. McClure, et al., Why has transparency evolved in aposematic butterflies? Insights from the largest radiation of aposematic butterflies, the Ithomiini. Proc. Biol. Sci. 286, 20182769 (2019).

35. E. Page, et al., Pervasive mimicry in flight behavior among aposematic butterflies. Proc. Natl. Acad. Sci. U. S. A. 121, e2300886121 (2024).

36. M. Elias, Evolution of aposematism and mimicry in butterflies: Causes, consequences and paradoxes. C. R. Biol. 342, 256–258 (2019).

37. T. N. Sherratt, The coevolution of warning signals. Proc. Biol. Sci. 269, 741–746 (2002).

38. K. R. Willmott, J. Mallet, Correlations between adult mimicry and larval host plants in ithomiine butterflies. Proc. Biol. Sci. 271 **Suppl 5**, S266–9 (2004).

39. N. Chazot, et al., Mutualistic mimicry and filtering by altitude shape the structure of Andean butterfly communities. Am. Nat. 183, 26–39 (2014).

40. D. Geale, Amateur’s Guide Butterflies Eastern Ecuador Peru (Willow Printing & Publishing Co, 2025).

41. P. E. Howse, Lepidopteran wing patterns and the evolution of satyric mimicry. Biol. J. Linn. Soc. Lond. 109, 203–214 (2013).

42. D. W. Kikuchi, D. W. Pfennig, Imperfect mimicry and the limits of natural selection. Q. Rev. Biol. 88, 297–315 (2013).

43. B. Rojas, et al., Multimodal aposematic signals and their emerging role in mate attraction. Front. Ecol. Evol. 6, 391900 (2018).

44. L. Maisonneuve, T. G. Aubier, The “sexual selection hypothesis” for the origin of aposematism. Evolution 79, 1153–1165 (2025).

45. J. Joshi, A. Prakash, K. Kunte, Evolutionary assembly of communities in butterfly mimicry rings. Am. Nat. 189, E58–E76 (2017).

46. S. B. Hagen, H. P. Leinaas, H. M. Lampe, Responses of great tits *Parus major* to small tortoiseshells *Aglais urticae* in feeding trials; evidence of aposematism. Ecol. Entomol. 28, 503–509 (2003).

47. M. Ilić, et al., Simple and complex, sexually dimorphic retinal mosaic of fritillary butterflies. Philos. Trans. R. Soc. Lond. B Biol. Sci. 377, 20210276 (2022).

48. A. Macdougall, M. Dawkins, Predator discrimination error and the benefits of Müllerian mimicry. Anim. Behav. 55, 1281–1288 (1998).

49. J. C. Oliver, K. A. Robertson, A. Monteiro, Accommodating natural and sexual selection in butterfly wing pattern evolution. Proc. Biol. Sci. 276, 2369–2375 (2009).

50. A. Vieira-Silva, G. B. Evora, A. V. L. Freitas, P. S. Oliveira, The relevance of flash coloration against avian predation in a Morpho butterfly: A field experiment in a tropical Rainforest. Ethology 130 (2024).

51. T. K. Suzuki, S. Tomita, H. Sezutsu, Gradual and contingent evolutionary emergence of leaf mimicry in butterfly wing patterns. BMC Evol. Biol. 14, 229 (2014).

52. T. K. Suzuki, S. Tomita, H. Sezutsu, Multicomponent structures in camouflage and mimicry in butterfly wing patterns. J. Morphol. 280, 149–166 (2019).

53. M. Stevens, I. C. Cuthill, A. M. M. Windsor, H. J. Walker, Disruptive contrast in animal camouflage. Proc. Biol. Sci. 273, 2433–2438 (2006).

54. D. J. Kemp, Female butterflies prefer males bearing bright iridescent ornamentation. Proc. Biol. Sci. 274, 1043–1047 (2007).

55. S. D. Finkbeiner, A. D. Briscoe, True UV color vision in a female butterfly with two UV opsins. J. Exp. Biol. 224, jeb242802 (2021).

56. S. D. Finkbeiner, D. A. Fishman, D. Osorio, A. D. Briscoe, Ultraviolet and yellow reflectance but not fluorescence is important for visual discrimination of conspecifics by *Heliconius erato*. J. Exp. Biol. 220, 1267–1276 (2017).

57. A. L. Klein, A. M. de Araújo, Sexual size dimorphism in the color pattern elements of two mimetic Heliconius butterflies. Neotrop. Entomol. 42, 600–606 (2013).

58. D. O. Rossato, et al., Subtle variation in size and shape of the whole forewing and the red band among co-mimics revealed by geometric morphometric analysis in Heliconius butterflies. Ecol. Evol. 8, 3280–3295 (2018).

59. K. S. Brown Jr, Geographical patterns of evolution in Neotropical Lepidoptera: differentiation of the species *Melinea* and *Mechanitis* (Nymphalidae, Ithomiinae). Syst. Entomol. 2, 161–197 (1977).

60. K. S. Brown Jr, J. Vasconcellos-Neto, Predation on Aposematic Ithomiine Butterflies by Tanagers (*Pipraeidea melanonota*). Biotropica 8, 136 (1976).

61. R. Deshmukh, S. Baral, A. Gandhimathi, M. Kuwalekar, K. Kunte, Mimicry in butterflies: co-option and a bag of magnificent developmental genetic tricks. Wiley Interdiscip. Rev. Dev. Biol. 7 (2018).

62. M. Joron, J. L. Mallet, Diversity in mimicry: paradox or paradigm? Trends Ecol. Evol. 13, 461–466 (1998).

63. J. Mallet, M. C. Singer, Individual selection, kin selection, and the shifting balance in the evolution of warning colours: the evidence from butterflies. Biol. J. Linn. Soc. Lond. 32, 337–350 (1987).

64. K. R. Willmott, J. C. Robinson Willmott, M. Elias, C. D. Jiggins, Maintaining mimicry diversity: optimal warning colour patterns differ among microhabitats in Amazonian clearwing butterflies. Proc. Biol. Sci. 284 (2017).

65. I. Birskis-Barros, A. V. L. Freitas, P. R. Guimarães Jr, Habitat generalist species constrain the diversity of mimicry rings in heterogeneous habitats. Sci. Rep. 11, 5072 (2021).

66. P. M. Brakefield, T. B. Larsen, The evolutionary significance of dry and wet season forms in some tropical butterflies. Biol. J. Linn. Soc. Lond. 22, 1–12 (1984).

67. L. P. Brower, J. Van Zandt Brower, Birds, butterflies, and plant Poisons: A study in ecological chemistry. Zoologica 49, 137–159 (1964).

68. P. R. Ackery, R. I. Vane-Wright, Milkweed butterflies (British Museum (Natural History), 1984).

69. K. S. Brown Jr, The biology of *Heliconius* and related genera. Annu. Rev. Entomol. 26, 427–457 (1981).

70. R. M. Merrill, et al., The diversification of *Heliconius* butterflies: what have we learned in 150 years? J. Evol. Biol. 28, 1417–1438 (2015).

71. N. Wahlberg, The phylogenetics and biochemistry of host-plant specialization in Melitaeine butterflies (Lepidoptera: Nymphalidae). Evolution 55, 522–537 (2001).

72. K. R. Willmott, The Genus Adelpha: Its Systematics, Biology and Biogeography (Lepidoptera: Nymphalidae: Limenitidini) (Scientific Publishers, 2003).

73. P. J. DeVries, Hostplant records and natural history notes on Costa Rican butterflies (Papilionidae, Pieridae & Nymphalidae). J. Res. Lepid. 24, 290–333 (1986).

74. N. Chazot, et al., Renewed diversification following Miocene landscape turnover in a Neotropical butterfly radiation. Glob. Ecol. Biogeogr. 28, 1118–1132 (2019).

75. R. I. Vane-Wright, On the definition of mimicry. Biol. J. Linn. Soc. Lond. 13, 1–6 (1980).

76. K. Kunte, Mimetic butterflies support Wallace’s model of sexual dimorphism. Proc. Biol. Sci. 275, 1617–1624 (2008).

77. M. Arias, et al., Transparency reduces predator detection in mimetic clearwing butterflies. Funct. Ecol. 33, 1110–1119 (2019).

78. J. J. Walker, Meetings of the Royal Entomological Society: Wednesday, December 1st, 1920. Trans. R. Entomol. Soc. Lond. 68, lxxxv–xcii (1921).

79. P. Eswaramoorthy, et al., Bokeh - Interactive Data Visualization in Python (2025).

80. G. Jocher, J. Qiu, Ultralytics YOLO11 (2024).

81. A. D. Warren, et al., Butterflies of America. Butterflies of America (2025). Available at: http://www.butterfliesofamerica.com/ [Accessed 10 July 2024].

82. M. Savela, Lepidoptera and some other life forms. Lepidoptera and some other life forms (2025). Available at: https://www.nic.funet.fi/pub/sci/bio/life/intro.html [Accessed 15 August 2024].

83. L. P. Brower, J. Van Zandt Brower, J. M. Corvino, Plant poisons in a terrestrial food chain. Proc. Natl. Acad. Sci. U. S. A. 57, 893–898 (1967).

84. L. P. Brower, D. O. Gibson, C. M. Moffitt, A. L. Panchen, Cardenolide content of *Danaus chrysippus* butterflies from three areas of East Africa. Biol. J. Linn. Soc. Lond. 10, 251–273 (1978).

85. P. Chai, Field observations and feeding experiments on the responses of rufous-tailed jacamars (*Galbula ruficauda*) to free-flying butterflies in a tropical rainforest. Biol. J. Linn. Soc. Lond. 29, 161–189 (1986).

86. F. Molleman, A. Kaasik, M. R. Whitaker, J. R. Carey, Partitioning variation in duration of ant feeding bouts can offer insights into the palatability of insects: experiments on African fruit-feeding butterflies. J. Res. Lepid. 45, 65–75 (2012).

87. F. Molleman, M. R. Whitaker, J. R. Carey, Rating palatability of butterflies by measuring ant feeding behavior. entomologische berichten 70 (2010).

88. J. R. Trigo, The chemistry of antipredator defense by secondary compounds in neotropical lepidoptera: facts, perspectives and caveats. J. Braz. Chem. Soc. 11, 551–561 (2000).

89. R. Nishida, “Chemical ecology of poisonous butterflies: Model or mimic? A paradox of sexual dimorphisms in müllerian mimicry” in Diversity and Evolution of Butterfly Wing Patterns, (Springer Singapore, 2017), pp. 205–220.

90. C. Pinheiro, Palatablility and escaping ability in Neotropical butterflies: tests with wild kingbirds (*Tyrannus melancholicus*, Tyrannidae). Biol. J. Linn. Soc. Lond. 59, 351–365 (1996).

91. A. Nahrstedt, R. H. Davis, Occurrence, variation and biosynthesis of the cyanogenic glucosides linamarin and lotaustralin in species of the Heliconiini (Insecta: Lepidoptera). Comp. Biochem. Physiol. B 75, 65–73 (1983).

92. R. B. Srygley, P. Chai, Predation and the elevation of thoracic temperature in brightly colored neotropical butterflies. Am. Nat. 135, 766–787 (1990).

93. R Core Team, R: A Language and Environment for Statistical Computing (R Foundation for Statistical Computing, 2024).

94. S. P. Blomberg, T. Garland Jr, A. R. Ives, Testing for phylogenetic signal in comparative data: behavioral traits are more labile. Evolution 57, 717–745 (2003).

95. S. N. Wood, Generalized Additive Models: An Introduction with R, Second Edition (CRC Press, 2017).

96. L. J. Revell, phytools 2.0: an updated R ecosystem for phylogenetic comparative methods (and other things). PeerJ 12, e16505 (2024).

